# Vascular calcification has a role in acute non-renal phosphate clearance

**DOI:** 10.1101/2020.07.29.225532

**Authors:** Mandy E Turner, Austin P Lansing, Paul S Jeronimo, Lok Hang Lee, Bruno A Svajger, Jason GE Zelt, Corey M Forster, Martin P Petkovich, Rachel M Holden, Michael A Adams

## Abstract

**Rationale:** Non-renal extravasation of phosphate from the circulation and transient accumulation into tissues and extracellular fluid is a regulated process of acute phosphate homeostasis that is not well understood. Following oral consumption of phosphate, circulating levels normalize long before urinary excretion has been completed. This process is especially relevant in the setting of chronic kidney disease (CKD), where phosphate exposure is prolonged due to inefficient kidney excretion. Furthermore, CKD-associated dysregulation of mineral metabolism exacerbates pathological accumulation of phosphate causing vascular calcification (VC).

**Objective:** Determine whether the systemic response to acute phosphate challenges is altered by the development and progression of VC.

**Methods/Results:** Acute circulating and tissue deposition of an acute phosphate challenge was assessed in two rat models of VC using radio-labelled phosphate tracer. In an adenine-induced model of CKD with VC, animals with VC had a blunted elevation of circulating ^33^PO_4_ following oral phosphate administration and the discordant deposition could be traced to the calcifying vasculature. In a non-CKD model of VC, VC was induced with 0.5ug/kg calcitriol and then withdrawn. The radio-labelled phosphate challenge was given to assess for vascular preference for phosphate uptake with and without the presence of an active calcification stimulus. The new transport to the calcifying vasculature correlates to the pre-existing burden of calcification, and can be substantially attenuated by removing the stimulus for calcification. The accrual is stimulated by a phosphate challenge, and not present in the same degree during passive disposition of circulating phosphate.

**Conclusions:** Our data indicate that calcifying arteries alter the systemic disposition of a phosphate challenge and acutely deposit substantial phosphate. This study supports the importance of diet as it relates to acute fluctuations of circulating phosphate and the importance of bioavailability and meal-to-meal management in CKD patients as a mediator of cardiovascular risk.

## Introduction

Medial vascular calcification (VC) is a pathology associated with aging, and is accelerated by diabetes and chronic kidney disease (CKD). Distinct from intimal calcification associated with atherosclerosis, in this pathology hydroxyapatite is actively deposited in the media and elastic lamina of muscular arteries. This pathology reduces vascular compliance and associates with poor cardiovascular outcomes^1,2^. Phosphate dysregulation has emerged as an important factor in the initiation and propagation of the calcification process. Serum phosphate, even in the upper ranges of normal, and at each stage of CKD, is recognized as an independent risk factor for cardiovascular disease^3^. Prevalence of VC in the thoracic aorta ranges from 37-60% in patients with stage 3 CKD when serum phosphate still within the normal range^4^.

Despite the growing recognition of circulating phosphate as a risk factor, less than 1% of total body phosphate content is found in the circulation. The tight regulation of circulating phosphate involves controlled movement within and between several compartments. These pools of phosphate typically include intestinal absorption of dietary phosphate, the movement of phosphate between skeletal, soft tissue and extracellular pools, and regulation of renal reabsorption and excretion. Phosphate transport in and out of these compartments is mediated, in part, through sodium phosphate-cotransporters (NaPi), as well as ubiquitous somatic phosphate inorganic transporters, PiT-1 and PiT-2. The activity and expression of NaPis are largely regulated by parathyroid hormone (PTH), fibroblast growth factor 23 (FGF-23) and calcitriol. Though not well understood, phosphate also undergoes paracellular transport along a concentration gradient in the intestine, an aspect of phosphate disposition which may be underestimated in CKD and potentially present in other tissues. Despite the clear role of phosphate in stimulating adaptive changes in its own regulation, the cellular mechanisms of phosphate-sensing remain poorly understood in somatic tissue^5^.

As kidney function declines, hormonal control mechanisms become unable to compensate and the resultant increase in circulating phosphate stresses cellular mechanisms of phosphate handling. The rise in circulating phosphate can occur acutely after a meal or, in later stages of CKD, present as chronic hyperphosphatemia. In a recent study using a rat model of CKD, the impact of oscillating from high to low dietary phosphate every two days resulted in VC much more severe than rats fed the same amount of phosphate without oscillations^6^. The burden of VC was comparable to that found in rats fed a continuously high dietary phosphate containing twice the overall amount of the oscillating burden. These findings suggest spikes in circulating phosphate may be an important driver of VC, potentially more important than overall exposure.

Our previous work indicates that acute responses to oral phosphate are already altered in mild to moderate CKD patients with normal serum phosphate^7^. Specifically, humans and rats with impaired kidney function but normal serum phosphate had a blunted elevation in their circulating phosphate following an oral phosphate challenge compared to those with health kidney function. Given that functional changes in phosphate absorption are not impaired in CKD^8^, this attenuated rise suggested that there were changes in the systemic distribution of the oral phosphate load in those with impaired kidney function, but did not provide evidence of the mechanism for this increased non-renal clearance.

There is little evidence for how a given tissue or organ is involved in the systemic disposition of phosphate following administration of an oral load, or how these processes are altered during the development of VC. In the present study, the objective was to determine whether the systemic response to acute phosphate challenges is altered by the development and progression of VC.

## Methods

All animal procedures were performed in accordance with the Canadian Council on Animal Care and were approved by Queen’s Animal Care Committee. Male Sprague Dawley rats (15-16 weeks, Hilltop Lab Animals Inc. PA, USA) were acclimated for a week prior to the start of the experiment and were individually housed and maintained on a 12-hour light/dark cycle throughout the duration of the study.

### Adenine-Induced CKD Model of Vascular Calcification

A chronic reduction in kidney function was induced using a 0.25% dietary adenine model for 5 weeks as previously described^9^ (Harland Teklad, TD.08672). A parallel control arm was ran concurrently without the dietary adenine, but otherwise identical diets (CON). After cessation of the adenine diet, animals were maintained on the non-adenine 0.5% phosphate diet for at least 4 days to ensure removal of the acute effects of dietary adenine (TD.150555), and then, at the sixth week, CKD and CON rats were stratified into high or low dietary phosphate according to bodyweight, circulating calcium and phosphate. The low dietary phosphate group remained on the 0.5% dietary formulation (LP) and the high dietary phosphate (HP) increased to 1% dietary phosphate (TD.08670) for two weeks. Blood was collected at least weekly from the saphenous vein. The total number of animals in each group are: CON-LP (N=13), CON-HP (N=11) CKD-LP (N=23) and CKD-HP (N=23).

### Administration of Oral Radiolabelled Phosphate

Two weeks following stratification into the dietary phosphate arms, animals were euthanized following an oral radiolabelled phosphate challenge. Animals were partially-fasted overnight to ensure consistency of stomach contents and then in the morning, animals were provided 2mL of sucralose gel (MediGel^®^, Clear H2O) with a total phosphate amount of 0.1g (equivalent to 100% daily intake of the LP animals, or 50% daily intake of HP animals). Phosphate in the gel was supplemented with dibasic and monobasic sodium phosphate salts (Sigma-Aldrich, Canada) and ~7.76 million Bq radio-labeled ^33^PO_4_ (NEN Radiochemicals). Animals were stratified by the three most recent measurements of serum creatinine, phosphate, calcium and bodyweight into one of three sacrifice times following the oral load of phosphate: 0 hour, 2 hours or 6 hours. Stratification metrics and final study animal numbers for each time point are outlined in Supplementary Table 1. Depending on sacrifice time, animals were sampled from the saphenous vein at 0, 20min, 40min, 1hr, 1.5hr, 2hr and then hourly until 6hr. Only two rats in the CKD-LP diet presented with VC and both animals were allocated *a priori* to the 6hr sacrifice time point. As a result, animals were excluded from analysis in Figures 2–3, as inclusion would have biased the vascular phosphate deposition findings for 6hr (but not 2hr) in CKD-LP animals. The rats sacrificed at 2 hours did not present a different profile than rats sacrificed at 6 hours (Supplementary Figure 1 and 2), as such Figure 2A-E presents the pooled combined profile of both groups at 0-2hr and statistics represent combined analysis.

### Non-CKD Calcitriol-Induced Model of Vascular Calcification

VC was induced through subcutaneous administration of suprapharmacological calcitriol (0.5μg/kg/day, Sterimax) for 8-days and maintained on a moderate 0.75% phosphate diet (Harland Teklad, TD.160324). A parallel control arm was completed concurrently. At day 7, animals (N=24) were stratified based on serum calcium, phosphate, PTH, FGF-23 and bodyweight into two time-points. The first group was sacrificed on experimental day 9, following 8 doses of calcitriol (Cx). The remaining rats no longer received calcitriol for 13 days (Post-Cx). Control animals were sacrificed at both time points and then combined due to no differences in any measurement. The number of animals in each group were: Cx (N=8), Post-Cx (N=9), Control Early (N=3), and Control Late (N=4). Blood was collected every 2-3 days via saphenous vein.

### IV Radiolabelled Phosphate Administration

Directly prior to sacrifice, animals were administered an intravenous load of radiolabelled phosphate. Intravenous delivery was chosen to bypass the potential effects of supraphysiologic calcitriol on gut phosphate transport. Under isoflurane anesthesia (2.5%, 2% O_2_), rats were administered 3mL of an isotonic sodium phosphate/sodium chloride solution containing 300μmol of phosphate and ~9.7 million Bq of ^33^PO_4_ (NEN Radiochemicals, Perkin Elmer) was infused intravenously into the jugular vein over 10 minutes (KD Scientific). Blood was sampled at baseline, 10 minutes, 20 minutes, and sacrifice (30 minutes).

A separate study was completed involving administering calcitriol subcutaneously via osmotic minipump (Alzet, 2mL capacity, 10μL/hr flow rate, 0.5μg/kg/day). Aside from method of administration, all other protocols were identical to the aforementioned first study. Under isoflurane anesthesia, the osmotic minipump was inserted on the back dorsolaterally, and subcutaneous meloxicam (2mg/kg loading, 1mg/kg maintenance) was administered pre- and post-operatively for 3 days. Animals were sacrificed 9 days after pump insertion. Animals were stratified into two groups based on serum calcium, phosphate, PTH, FGF-23 and bodyweight at day 7. One group (N=6) received the intravenous infusion of a 300μmol phosphate spiked with radiolabeled phosphate as described above. The second group (N=6) received an infusion of only the tracer amount of radiolabeled phosphate in saline, but lacking the phosphate load.

### Tissue Harvest and Tissue Assessment Preparation

Animals were anesthetized with isoflurane (5%) and sacrificed via cardiac puncture and exsanguination. Urine was collected directly from the bladder. Gastrointestinal tissue from the stomach to anus was quickly excised and separated. Samples of chyme were collected from the stomach, proximal small intestine (duodenum) and distal small intestine (ileum) and large intestine. Feces was collected from the distal colon. Multiple somatic tissue types were collected (n=50) including various samples of arteries, veins, cardiac and skeletal muscles, bone, kidney, fat, intestine, liver, pancreas, and lung. Tissues and chyme were demineralized in 1N HCl for 1 week and minerals and radioactivity were measured in the acid homogenate.

### Biochemical blood and urinary measurements

Serum creatinine as well as both serum and urinary calcium and phosphate were evaluated spectrophotometrically (SynergyHT Microplate Reader). Creatinine was evaluated using the Jaffe method (QuantiChrom Creatinine Assay Kit, Bioassay Systems). Serum and tissue calcium was measured using the o-cresolphthalein method^10^ and free phosphate was measured using the malachite green (Sigma-Aldrich) method as described by Heresztyn and Nicholson^11^. Plasma levels of intact PTH and C-terminal/intact FGF-23 were measured by ELISA (Immunotopics Inc.).

### Radioactivity measurement and analysis

For radioactivity assessments, tissue acid homogenates, serum, and urine samples were added to Ultima Gold AB scintillation cocktail (Perkin Elmer) and analyzed using a Beckman Coulter LS 6500 multi-purpose scintillation counter. Each sample was measured twice for a 1-minute count time. Corrected radioactivity was obtained by subtracting background from all samples and then normalized to the amount of radioactivity ingested by each rat. Serum specific activity was calculated at each time point over the course of the study (equation 1). In order to transform counts/mg of tissue to an estimation of amount of phosphate accrued per tissue, a time-weighted average serum specific activity was generated and was used to estimate tissue phosphate accrual (equation 2) as described previously^12^.

1. Serum Specific Activity (μCi/pmol) = Serum Radioactivity (μCi/uL) / Serum Phosphate (mM)
2. Tissue PO_4_ Accrual (pmol PO4/mg tissue) = Tissue Radioactivity (μCi/mg tissue) / Average Specific Activity (μCi/pmol)

### Von Kossa Histology

The arteries were fixed in 10X neutral phosphate-buffered saline with 4% paraformaldehyde and embedded in paraffin blocks. Sections (4 μm) were stained for calcification using the Von Kossa method as previously described^13^. Areas of calcification appeared as dark brown regions in the medial wall of the artery.

### Analysis

Text data is represented as mean±SD, unless otherwise indicated. The threshold for significance was a p-value <0.05. All statistical tests and graph generation were done on GraphPad Prism (Version 8.4). Statistics performed are outlined in detail in figure captions and table footnotes.

## Results

The dietary-adenine model of CKD was confirmed by elevated serum creatinine (Table 1). CKD rats also had elevated serum phosphate, PTH, and FGF-23 that was exacerbated by the addition of high dietary phosphate (CKD-HP), compared to controls. A chronic increase in dietary phosphate did not significantly alter any of the measured parameters in control animals. Assessments of weekly increases in serum creatinine and phosphate are presented in Supplementary Figure 3.

**Table 1:** CKD-MBD Rat Model Characteristics at Sacrifice

| Treatment Group | Creatinine (μM) | PO<sub>4</sub> (mM) | Ca (mM) | PTH (pg/mL) | FGF23 (pg/mL) | Bodyweight (g) |
| --- | --- | --- | --- | --- | --- | --- |
| **CKD-LP** | 441.4±100.0<br>4a, 4b | 3.74±0.80<br>2a, 4b | 2.79±0.52 | 930 [542,1479]<br>4a, 4b | 4192 [3322,7226]<br>4a, 4b | 465±27<br>3a, 1b |
| **CKD-HP** | 512.4±166.6<br>4a, 4b | 5.05±0.98<br>4a, 4b, 4c | 2.47±0.72 | 3100 [2497,6090]<br>4a, 4b, 4c | 68176 [47507,85767]<br>4a, 4b, 4c | 411±25<br>4a, 4b, 4c |
| **CON-LP** | 41.7±8.8 | 2.47±0.62 | 2.51±0.28 | 131 [99, 220] | 326[306,383] | 506±22 |
| **CON-HP** | 42.8±10.4 | 2.04±0.23 | 2.57±0.31 | 243 [109, 296] | 306[287,320] | 497±28 |
| *CKD Intervention* | *** | *** | ns | *** | *** | *** |
| *Dietary Phosphate* | ns | *** | ns | *** | *** | *** |
| *Interaction* | ns | *** | ns | *** | *** | *** |

Two-way ANOVA with Sidak's multiple comparisons test. <sup>a</sup>Difference from CON-LP <sup>b</sup>Difference from CON-HP <sup>c</sup>Difference from CKD-LP. PTH. 1b: p<0.05, 2a: p<0.01. 3a: p<0.001, 4a or 4b or 4c: p<0.0001. PTH/FGF-23 represented as median [IQR] and statistics performed on normalized log-transformed data. Values for phosphate, calcium, PTH reflect directly prior to oral phosphate load. Significant sources of variation as identified by two-way ANOVA bottom three rows. \*\*\*p<0.001

High dietary phosphate in CKD animals induced consistent medial layer vascular calcification (VC), as indicated by substantial elevations of calcium and phosphate in both central (22/23; 96%) and distal arteries (23/23, 100%). This finding was confirmed histologically using von Kossa staining (Figure 1A-C). The rats fed low phosphate (CKD-LP) were not significantly different from controls, with only two rats (2/23, 8.7%) developing detectable VC. Taken together with circulating markers, the high dietary-phosphate group with adenine-induced CKD had changes characteristic of CKD-MBD.

**Figure 1:**
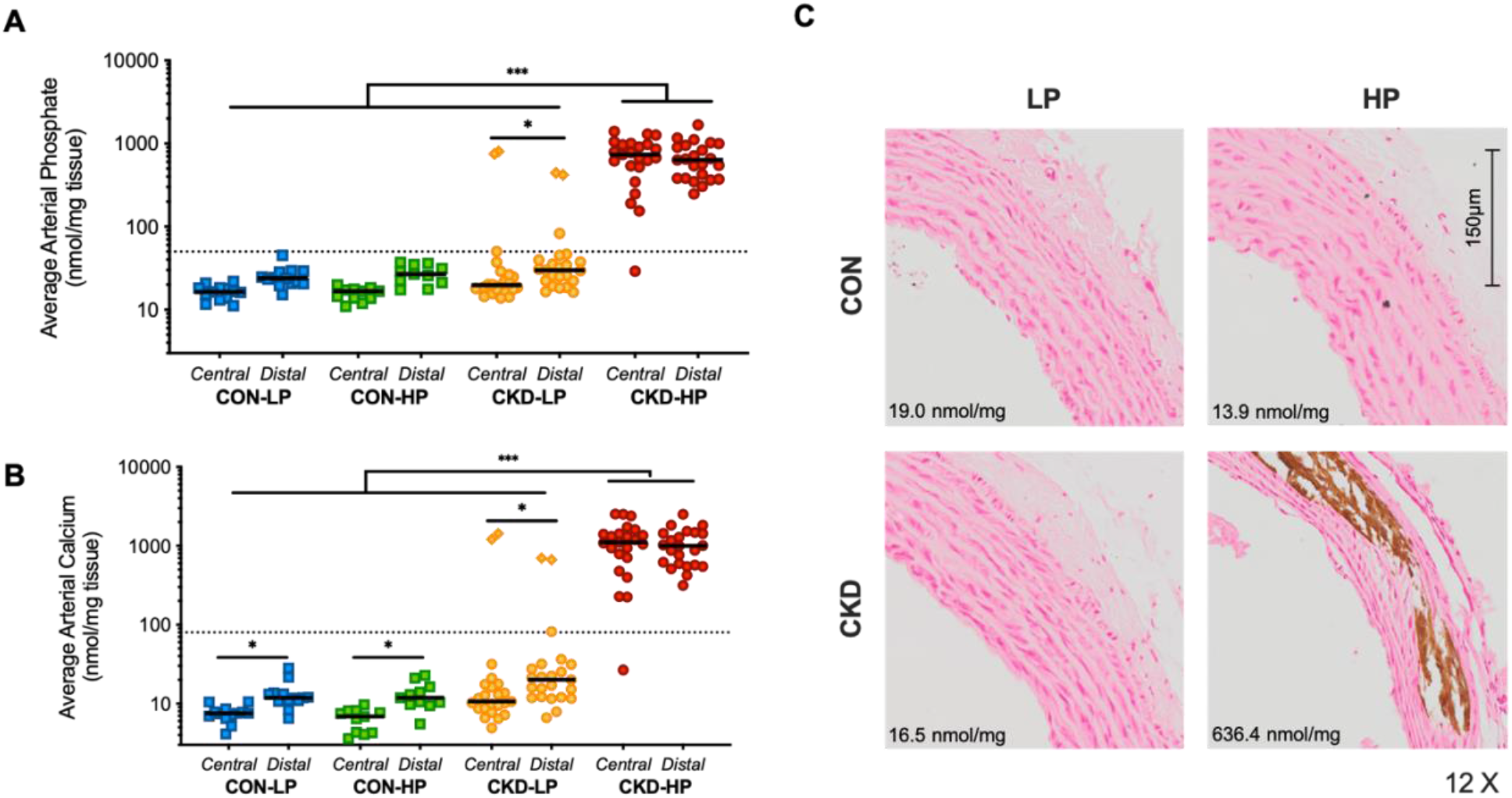
Increased dietary phosphate induces medial vascular calcification in arterial tissue in experimental CKD. **(A)** Arterial phosphate and **(B)** calcium per mg of wet weight tissue. Line at median. Each data point represents the mean of central (N=5) and peripheral (N=10) vessels in each rat. Dotted line at 50 ng/mg tissue phosphate and 80 ng/mg tissue calcium, the approximate mineral levels needed to detect vascular calcification histologically via von Kossa staining. Three-way ANOVA on log-transformed data with *post hoc* Tukey-corrected multiple comparisons. *p<0.05, **p<0.01 *** p<0.001. **(C)** Representative visible aortic medial calcification indicated by von Kossa phosphate staining in only CKD-HP. Tissue phosphate indicated on image.

In response to the oral phosphate load, total serum phosphate increased acutely in all groups, although only significantly in CKD, which occurred at 1hr and remained elevated for the remainder of the 6 hr analysis. At all points, total circulating phosphate is higher in CKD-HP than CKD-LP (Figure 2A), however, the chronic dietary phosphate did not impact the magnitude of the absolute change in circulating phosphate at any time points (Figure 2B). In contrast, the chronic change in dietary phosphate altered the responsiveness of PTH to the acute oral phosphate load. That is, only in the rats on low phosphate diet did the PTH rise significantly from baseline in response to the acute phosphate load at 1 hour (Figure 2C-D). Circulating FGF-23 was not significantly increased by the acute phosphate load (Supplementary Figure 4). There were minimal changes in serum calcium at the measured time points (Supplementary Figures 1, 2).

**Figure 2:**
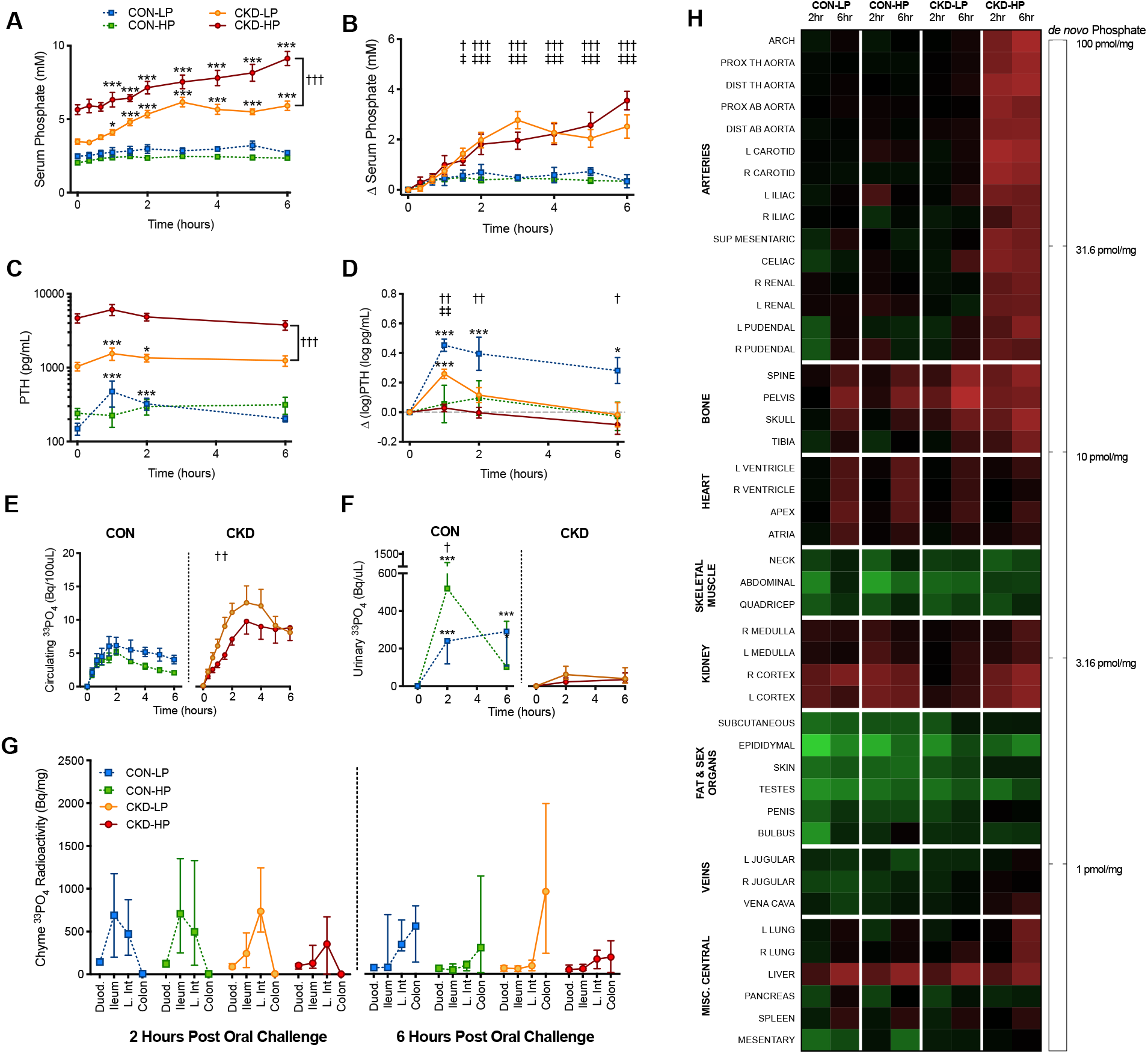
Acute response and tissue deposition to radiolabeled phosphate challenge altered by dietary phosphate in experimental model of CKD. **(A)** Circulating total phosphate, **(B)** absolute change in total phosphate, **(C)** circulating PTH, **(D)** absolute change in log-transformed PTH, **(E)** circulating ^33^PO_4_ per 100uL of serum, and **(F)** urinary ^33^PO_4_ per uL. Repeated measures mixed effects model analysis with *post hoc* tests evaluating within-group differences from time 0 (Dunnett’s correction; * p<0.05, *p<0.01, ***p<0.001) and between group differences comparing CKD-LP to CON-LP (†) and CKD-HP to CON-HP (‡) at each time point, unless indicated otherwise (Tukey correction; † p<0.05, ††† p<0.001). Data expressed as mean ± SEM. Comparisons of PTH panel C, evaluated on log-transformed data. **(G)** Change in the profile of radioactive phosphate along the gastrointestinal tract at 2 hours and 6 hours following an oral load of radioactive phosphate. Radiolabeled phosphate load along intestinal tract indicates at 2 hours following oral load, small intestinal absorption is still occurring, and has finished by 6 hours in all groups. **(H)** Experimental CKD and dietary phosphate alter tissue disposition of an oral load of radiolabeled phosphate. *De novo* tissue phosphate accrual in various tissues across the body at 2 and 6 hours following oral load grouped by tissue type. The heat map coloration represents the amount of *de novo* phosphate accumulated per mg of wet-weight tissue on a logarithmically-transformed scale. Arteries sorted by external diameter. For full tissue list see supplementary methods.

Consistent with declining kidney function, circulating ^33^PO_4_ elevated more in the CKD rats than in the controls (Figure 2E, statistics not shown). However, there was also significant impact (p<0.05, Three-way ANOVA) of chronically increasing the dietary phosphate on the circulating ^33^PO_4_. Specifically, there was a blunted elevation of circulating ^33^PO_4_ in the CKD-HP group at 1.5 and 2 hours.

As expected, renal phosphate clearance was decreased in CKD rats compared to controls (Figure 2F). Control animals with consuming increased dietary phosphate appeared to have more robust urinary excretion of phosphate, as measured by ^33^PO_4_. There is no evidence of altered calcium excretion following the oral load of phosphate in CKD animals (Supplementary Figure 5).

Chyme radioactivity was used as a marker of the absorption/intestinal excretion profile of the acute phosphate load. There was no measurable impact of CKD or the chronic dietary phosphate on this profile (Figure 2G, Three-way ANOVA). In all groups, at 2 hours, there is significantly higher amount of ^33^PO_4_ in the chyme and small intestine and very little in the feces. The opposite is true at 6 hours, at which time there is significant fecal ^33^PO_4_. The 2-hour time point reflects the status during absorption and the 6-hour time point reflects the status after most intestinal absorption has already occurred.

Figure 2H depicts estimated amount of *de novo* phosphate across all tissues at 2 and 6 hours in each treatment group. Tissues were grouped according to function and/or location. Supplementary Figure 7 is a grey-scale depiction of the heat map. Across all treatment groups and dietary phosphate interventions, the most substantial localization occurred in the bone, kidneys, liver and cardiac muscle.

At both 2 and 6 hours following the oral load, there was a significant impact of dietary phosphate on the *de novo* phosphate accrual in the arteries, whereby accrual was markedly elevated in CKD animals fed a high phosphate diet, compared to those fed a low phosphate diet or control (Figure 3A). This finding was consistent throughout the vascular tree. The accrual in the vasculature of the CKD animals fed a low phosphate diet, which were uncalcified, was similar to that of the controls. There was no difference in accrual between 2 and 6 hours in any of the treatment groups.

**Figure 3:**
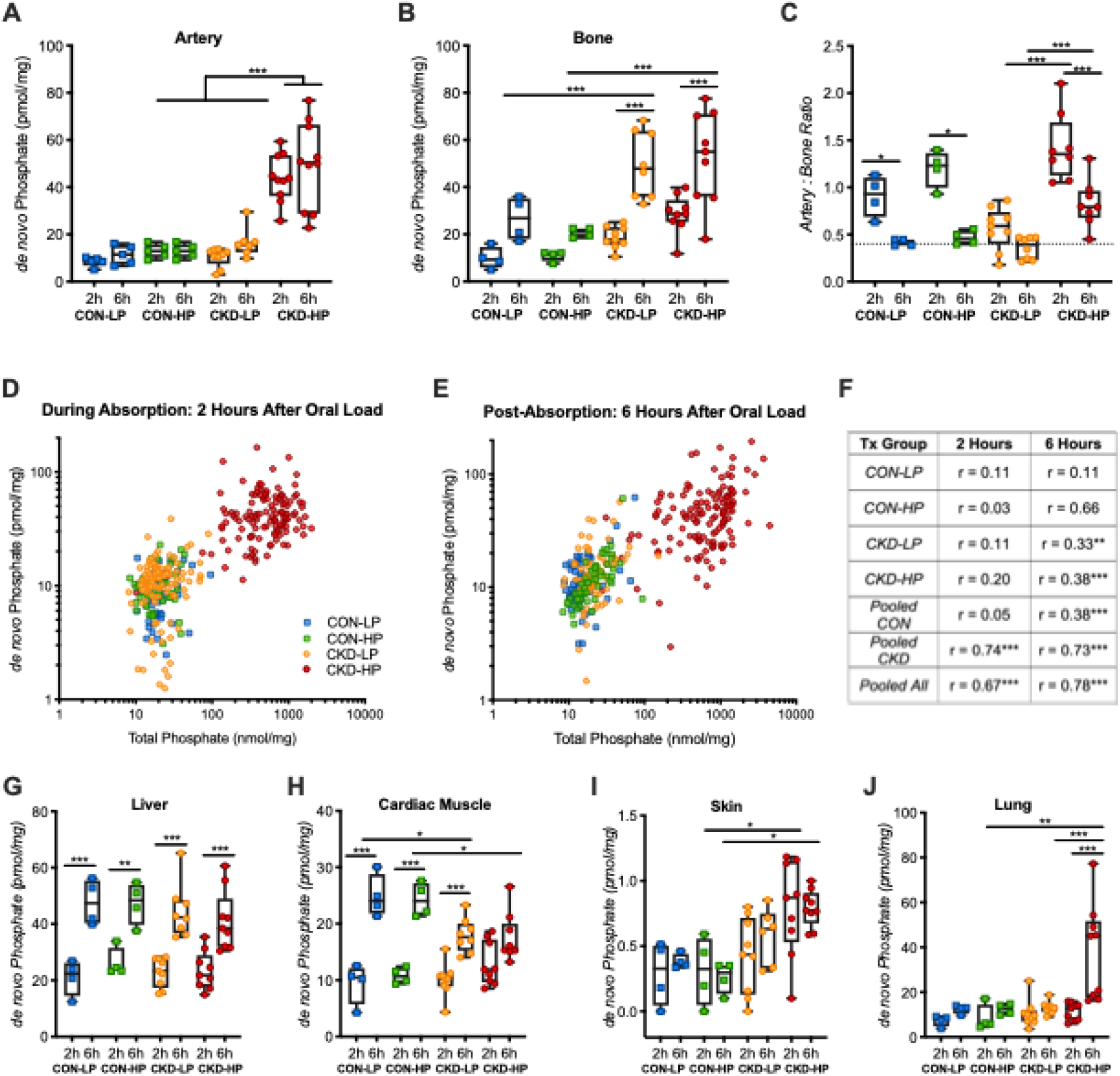
Calcified vascular tissue is a depot of *de novo* phosphate in the setting of experimental CKD and high dietary phosphate. Comparison of *de novo* phosphate deposition in **(A)** arteries**, (B)** bone, and **(C)** the ratio artery:bone at 2 and 6 hours after the oral phosphate. Each data point represents the pooled average of the 15 arterial or 4 bone samples for each individual rat. Dotted line on pane C represents pooled control means at 6 hours. Three-way ANOVA with *post hoc* Sidak-corrected multiple comparisons. *p<0.05, **p<0.01 *** p<0.001 between data-sets that differed only by one variable. (A) CKD intervention (p<0.001), dietary phosphate (p<0.001), and their interaction (p<0.05) were significant sources of variation. (B) CKD intervention (p<0.001), time of sacrifice (p<0.001), and their interaction (p<0.05), but not dietary phosphate, were significant sources of variation. (C) The interaction of dietary phosphate and CKD intervention (p<0.001), time of sacrifice (p<0.001), dietary phosphate (p<0.001), and their interaction (p<0.05) were significant sources of variation. **(D-F)** *De novo* phosphate accumulation correlates with present vascular calcification. Change in *de novo* arterial phosphate accumulation as a function of total tissue phosphate at 2 hours (D) and 6 hours (E) following the oral load. Each data point is from a single artery sample (15 different artery samples x 51 animals = 795 data points). Spearman correlation r-values of each group and pooled groups (F). *p<0.05, **p<0.01, ****p<0.001. Comparison of *de novo* phosphate deposition in **(G)** liver, **(H)** cardiac muscle, **(I)** skin, and **(J)** lung 2 and 6 hours after the oral phosphate.

In contrast, while there was no impact of dietary phosphate on the *de novo* accrual phosphate in the bone, there was an impact of CKD treatment and time of sacrifice (Figure 3B). Specifically, in each group there was more accrual at 6 hours than at 2 hours, which was exacerbated by CKD, likely a result of reduced clearance capacity. The arterial-to-bone accrual ratio exceeds 1 in CKD-HP, indicating the accrual in the vessels per mg of tissue is higher than that of bone, and at 6 hours normalizes to 1 (Figure 3C).

In CKD animals, *de novo* accrual into the vessels correlates strongly with the resident tissue phosphate as an indicator of VC at 2 and 6 hours (r > 0.67, p<0.0001) (Figure 3D-F). At 2 hours within each treatment group, a phase during which absorption is occurring, the correlation is weak and non-significant. However, at 6 hours following absorption, the correlation strengthens in each group, and there is a strong relationship between *de novo* phosphate and the total tissue phosphate in all groups, except CON-LP.

Liver accrual of phosphate was not impacted by CKD or dietary phosphate, but skin and lung had substantially more accrual in CKD-HP (Figure 3GIJ). Cardiac muscle, however, had attenuated phosphate uptake in CKD at 6hrs (Figure 3H)

In a non-CKD model of medial VC, administration of 0.5 μg/kg calcitriol for 8 days resulted in hypercalcemia, transient hyperphosphatemia, suppression of PTH, and marked elevation in FGF-23 (Figure 4A-D). In the subset of animals sacrificed while the stimulus was still present, 8 days of calcitriol (Cx) was sufficient to generate substantial medial VC, as indicated by elevations in arterial calcium and phosphate (Figure 4E-F) and confirmed histologically by Von Kossa staining (Figure 4G). Thirteen days after the cessation of stimulus (Post-Cx), circulating parameters of mineral metabolism had normalized, however calcification was non-reduced and histologically similar to that from animals sacrificed earlier.

**Figure 4:**
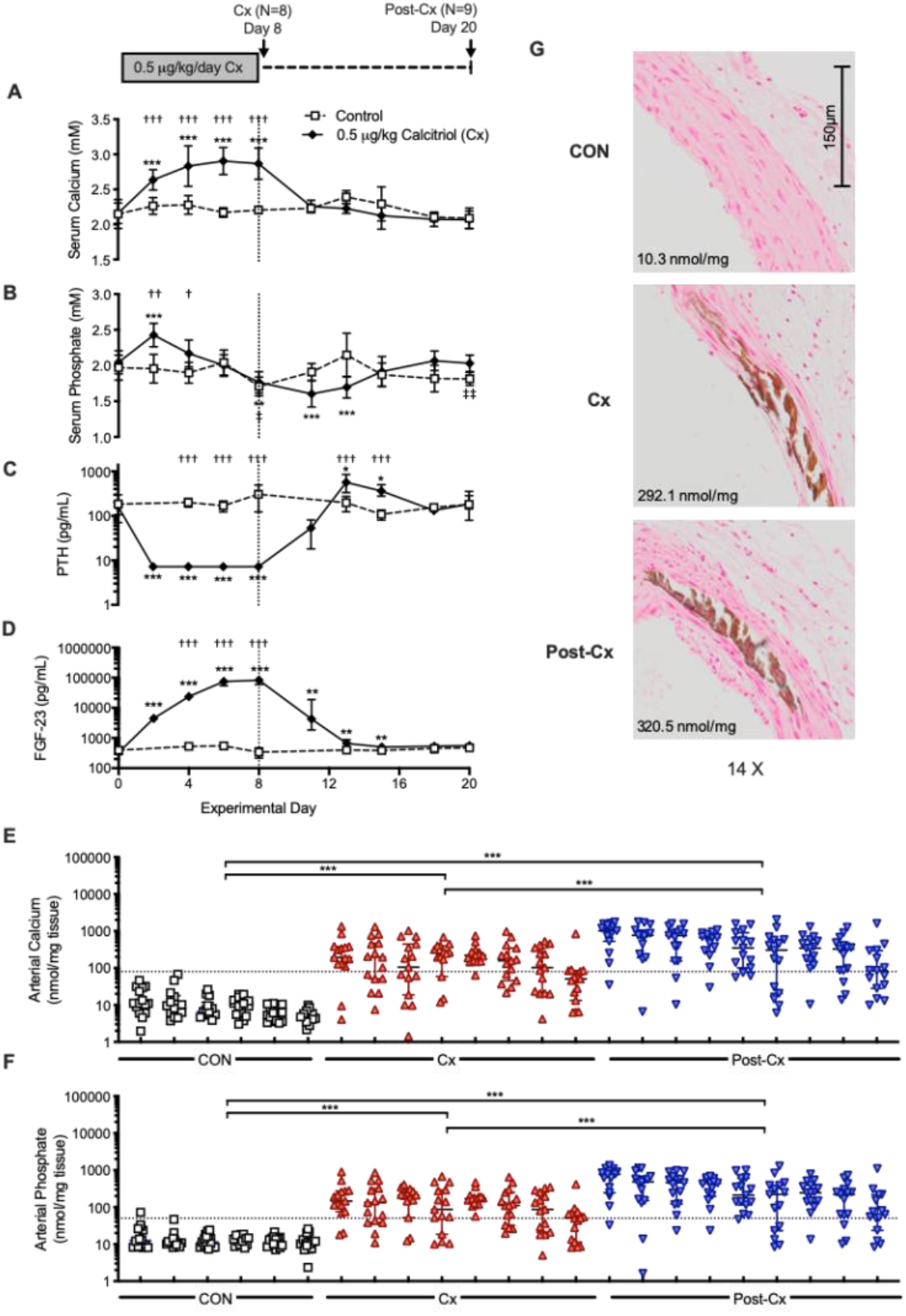
Persistence of vascular calcification in non-CKD model after removal of the calcification stimulus and normalization of circulating markers. Perturbations of circulating **(A)** calcium, **(B)** phosphate, **(C)** PTH, **(D)** FGF-23 following dosage of calcitriol (0.5 μg/kg/day) for 8 rats (Cx). After cessation of treatment for 12 days, circulating parameters largely returned to normal (Post-Cx). Repeated measures mixed effects model analysis with *post hoc* tests evaluating within-group differences from time 0 (Dunnett’s correction; * p<0.05, *p<0.01, ***p<0.001) and between group differences comparing Cx to Post-Cx (†) (Tukey correction; † p<0.05, ††† p<0.001). Data expressed as mean ± SD (A-B) or median IQR (C-D). Comparisons of PTH and FGF-23 evaluated on log-transformed data. Persistent vascular calcification, as indicated by **(E)** arterial calcium and **(F)** phosphate after removal of stimulus. Each column represents an animal, and each data-point a vascular tissue measured. Dotted line at 50 ng/mg tissue phosphate and 80 ng/mg tissue calcium, the mineral levels needed to detect vascular calcification histologically via von Kossa staining. Two-way ANOVA on log-transformed data with *post hoc* Tukey-corrected multiple comparisons. ***p<0.001. **(G)** Representative visible medial calcification indicated by Von Kossa phosphate staining in both the Cx and Post-Cx rats.

In response to a 10-minute intravenous infusion of phosphate with tracer ^33^PO_4_, serum phosphate elevated similarly in all groups (Figure 5A), and there was a reduction in calcium in all groups by 20 minutes (Figure 5B). At all time-points, serum calcium was higher in the Cx group, and in response to the IV phosphate, some animals had a substantial elevation in calcium at 10 minutes (Figure 5B). Cx animals had a blunted elevation in ^33^PO_4_ compared to controls and Post-Cx animals (Figure 5C).

**Figure 5:**
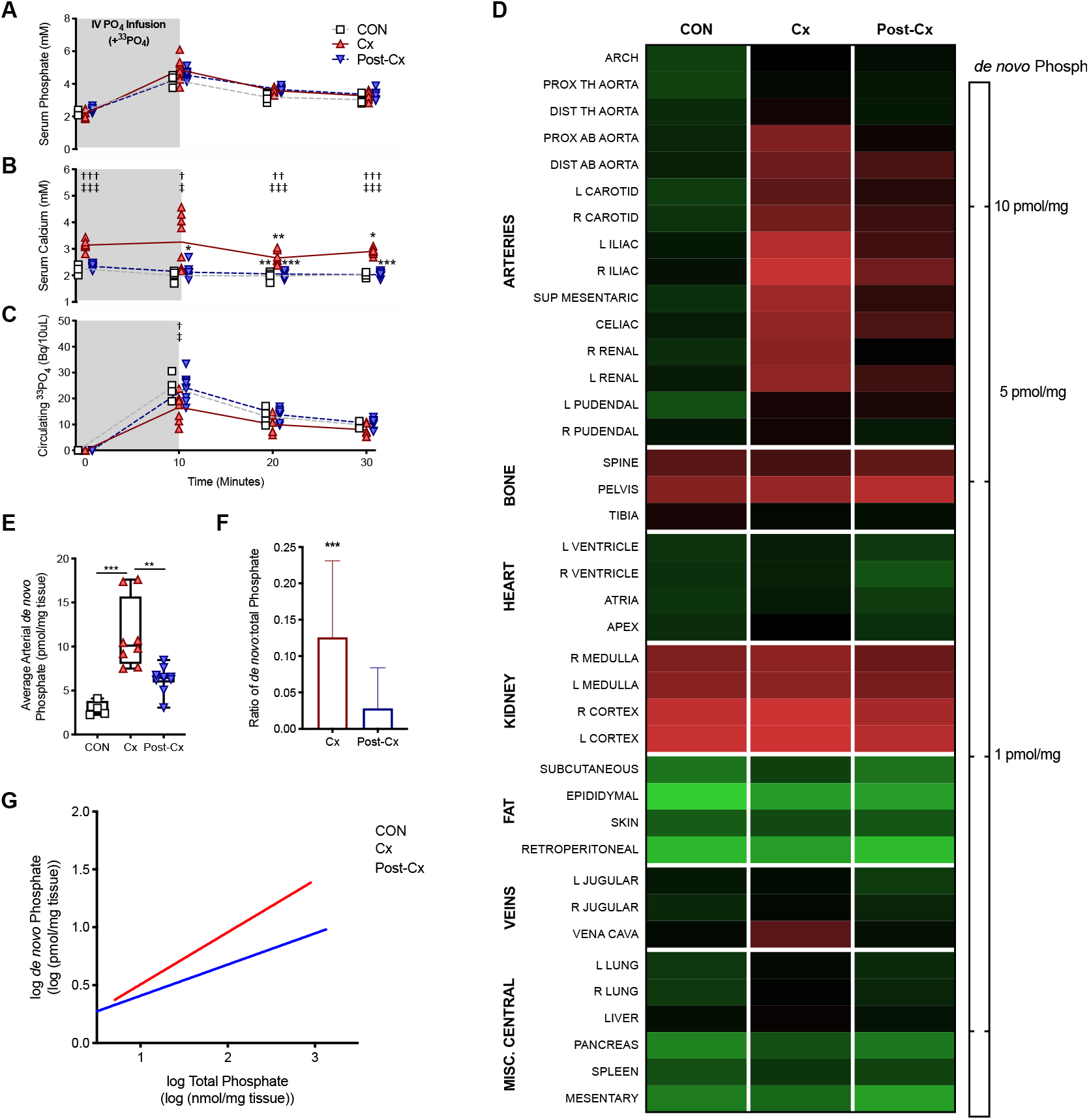
Presence of a stimulus for calcification differentially impacts acute deposition in calcified vessels. **(A)** Circulating phosphate, **(B)** calcium, and **(C)** radiolabeled ^33^PO_4_ during and after the IV infusion of a radiolabeled phosphate load. Repeated measures two-way ANOVA with *post hoc* tests comparing Cx – CON(†), and Post-Cx to CON (‡) (Tukey correction; † p<0.05, ††† p<0.001) and evaluating within-group differences from time 0 (Dunnett’s correction; * p<0.05, *p<0.01, ***p<0.001). At no time point was Post-Cx different than CON. At all time-points, serum phosphate and circulating ^33^PO_4_ was higher than at baseline. Line at mean. **(D)** *De novo* tissue phosphate accrual in various tissues across the body grouped by tissue type. The heat map coloration represents the amount of *de novo* phosphate accumulated per mg of wet-weight tissue on a logarithmically-transformed scale. Arteries sorted by external diameter. For full tissue list see supplementary methods. **(E)** Average arterial *de novo* phosphate for each animal. Kruskal-Wallis with *post hoc* Dunn’s multiple comparison test (**p<0.01, ***p<0.001). **(F)** Ratio of *de novo* arterial phosphate to total arterial phosphate for each vessel. Median, IQR, 95% CI shown. Data points outside 95% CI plotted. Mann-Whitney test (**p<0.01). *De novo* phosphate accumulation in arteries plotted against total tissue phosphate following IV infusion. Each data point is from a single artery sample. Log-log linear regression plotted (Cx: R^2^=0.58, y=0.45x+0.05, Post-Cx: R^2^=0.40, y=0.27x+0.14).

Estimated tissue accrual of phosphate in response to the phosphate challenge is presented in Figure 5D. Supplementary Figure 8 is a greyscale depiction. In a mixed effects model, each tissue was compared between treatment groups (statistics not presented on figure, Supplementary Table 2). There was no difference between the groups in acute accrual in most tissue groups, specifically bones, kidney, adipose and veins. However, there was a substantial difference in average arterial accrual, in that calcified animals currently receiving stimulus (Cx) had ~4X more deposition than controls, and control and calcified animals after the calcification stimulus was removed (Post-Cx) were not different from each other (Figure 4E). Similarly, there was a correlation between VC and *de novo* phosphate in both calcified groups, however there was substantially more phosphate accrual for a given amount of calcification in the Cx group with the mean ratio lower by ~80% (Figure 5F-G).

In order to elucidate the role of a phosphate challenge, as opposed to non-challenged movement of circulating phosphate, the IV radiolabeled phosphate infusion was compared to a saline infusion spiked with ^33^PO_4_ (Figure 6A-B). During the time-frame of the IV challenge, the phosphate load stimulated urinary excretion of phosphate (Figure 6C). Subtracting the mean accrual of each tissue in the saline group from the phosphate group, we generate the differential accrual as a result of the phosphate challenge (Figure 6D). As expected, the kidneys exhibited the greatest difference, followed by bones, and then central and peripheral arteries, and vein. In order to assess the impact of pre-existing VC on the accrual, we see that in the setting of no-, or mild-VC, the accrual is similar regardless of whether or not there was a phosphate challenge was applied, however as calcification progressed, the phosphate challenge preferentially deposited in the vasculature. (Figure 6E-F).

**Figure 6:**
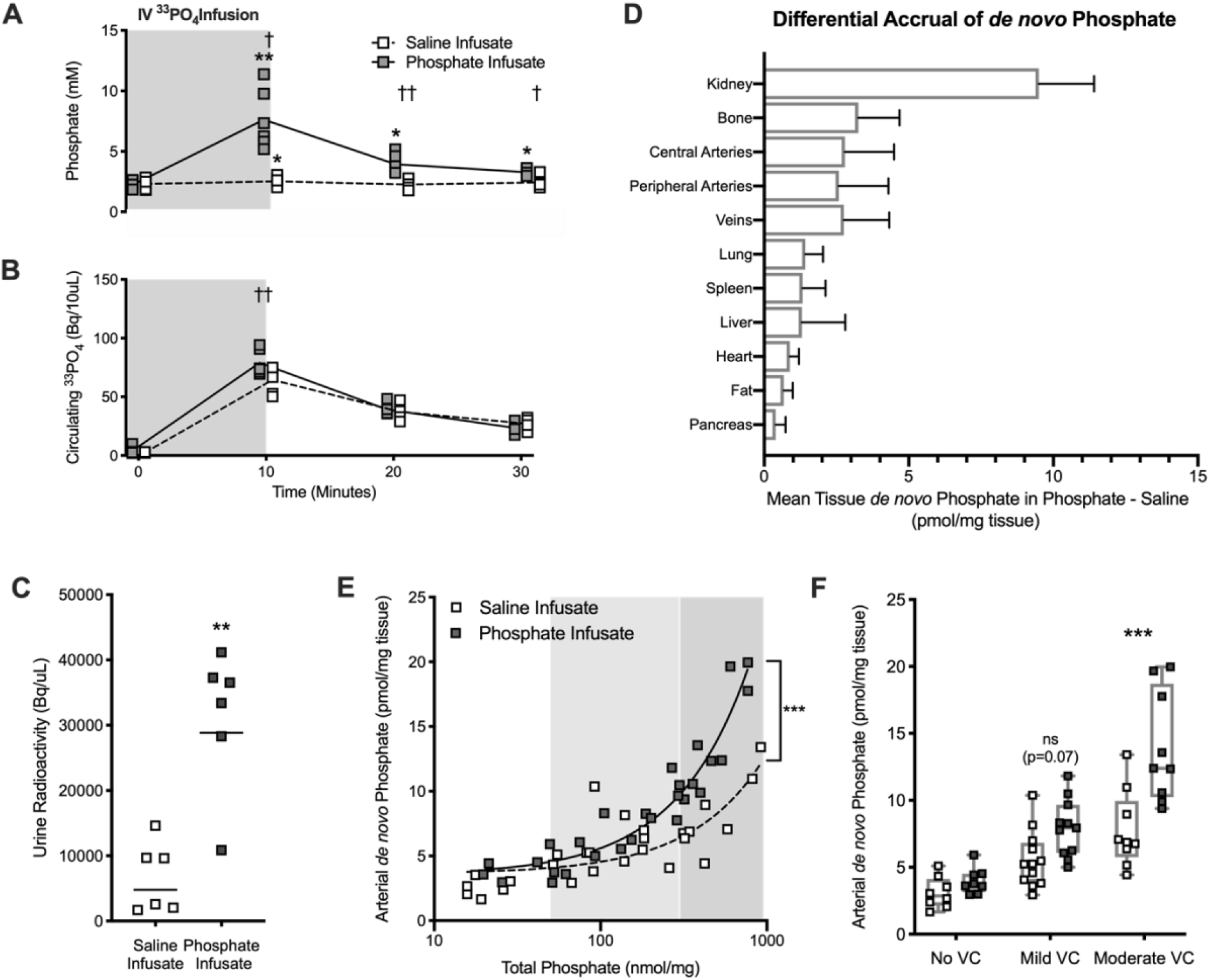
Acute phosphate acts as a stimulus for differential vascular deposition in calcified vessels only. **(A)** Circulating phosphate and **(B)** radiolabeled ^33^PO_4_ during and after the IV infusion of either a radiolabeled phosphate load or saline with tracer. Repeated measures two-way ANOVA with *post hoc* tests comparing within-group differences from time 0 (Dunnett’s correction; * p<0.05, *p<0.01, ***p<0.001) and between infusates at each time point (Sidak correction; † p<0.05, ††† p<0.001). Line at mean. **(C)** Urine radioactivity compared using Mann-Whitney test, line at geometric mean. **(D)** Differential *de novo* phosphate accrual in phosphate versus saline infusion. Mean tissue *de novo* phosphate in saline infusion group subtracted from phosphate group. Mean SD. **(E)** *De novo* phosphate accumulation in arteries plotted against total tissue phosphate following IV infusion. Each data point is from a single central artery sample. Linear regression plotted (Phosphate: R^2^=0.91, y=0.021x+3.5, Saline: R^2^=0.60, y=0.009x+3.6). Slopes of lines are significantly different p<0.001. **(F)** Calcified vasculature selectively accrues more phosphate acutely. Degree of calcification binned according to no VC (<50nmol/mg), mild VC (50-300nmol/mg), and moderate VC (>300 nmol/mg). Two-way ANOVA with *post hoc* Sidak-corrected comparisons between infusates (**p<0.01).

## Discussion

In the present study, we demonstrate that calcifying vasculature alters the systemic disposition of a phosphate challenge by acutely depositing phosphate during the process of tissue and extracellular equilibration thereby buffering circulating phosphate (Figure 7). Using a radiolabelled oral phosphate challenge in a rat model of CKD-MBD, the studies revealed a blunted rise in circulating ^33^PO_4_, that associated with differential tissue deposition towards the calcifying vasculature. We confirmed that tissue accrual was stimulated by an acute phosphate challenge, given that the same level of deposition did not occur solely with passive disposition of circulating phosphate. The extent of new transport to the calcifying vasculature correlates to the pre-existing burden of calcification, and can be substantially attenuated by removing the stimulus for calcification, indicating it is not a by-product of high hydroxyapatite burden, but the consequence of an active calcification process.

**Figure 7:**
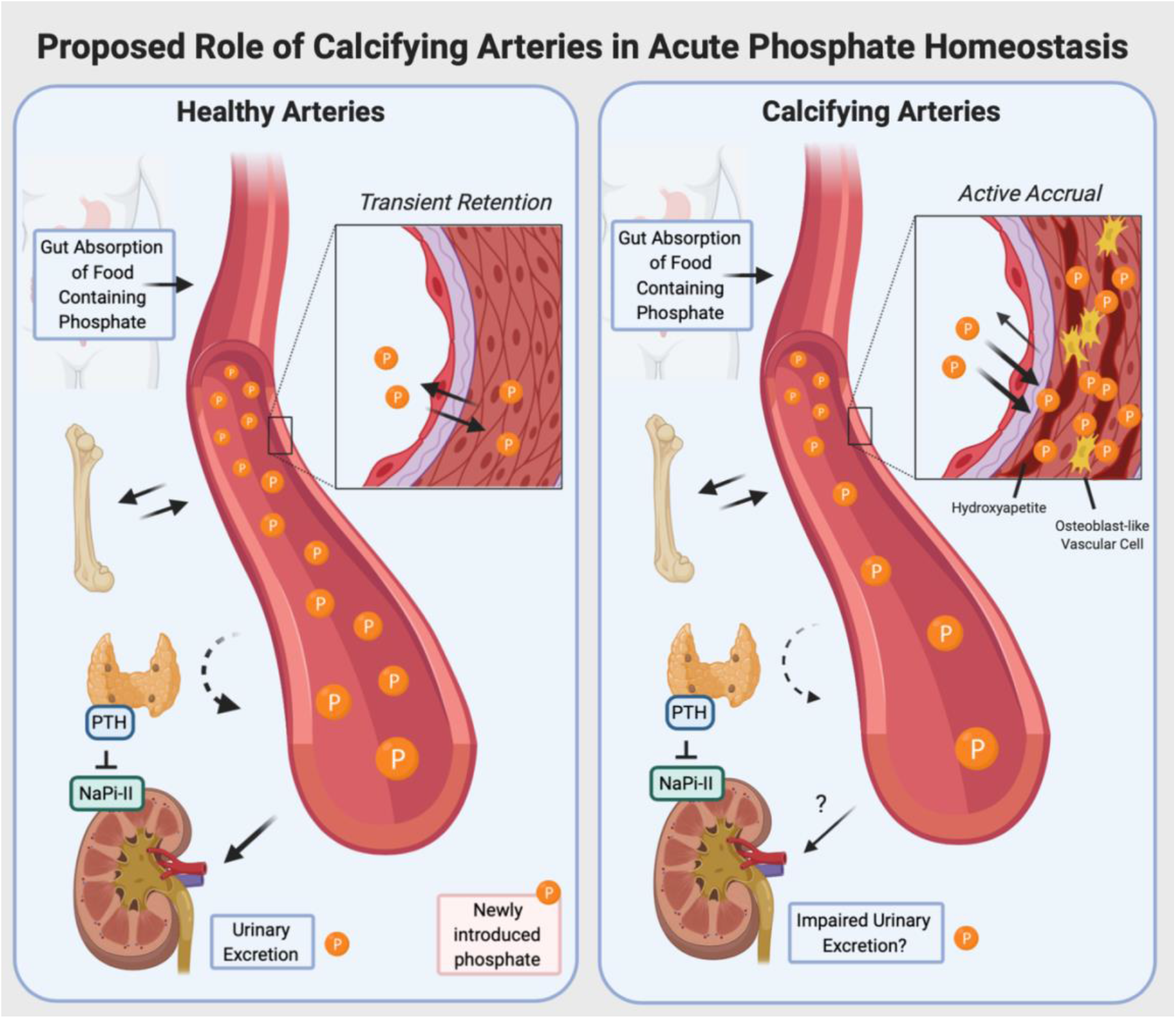
Proposed conceptual framework describing the role of calcifying vasculature in the response to an acute challenge of oral phosphate (i.e. a meal) and consequences. In the setting of actively calcifying vasculature, elevations in phosphate following an oral absorption of a phosphate containing food item, phosphate is preferentially removed from the circulation and into calcifying vasculature. This blunting of circulating phosphate may have impacts on other aspects of acute hormonal phosphate response (i.e. impairing PTH signalling) and impair timely urinary excretion.

The findings from this study indicate that non-renal clearance of phosphate from the circulation resulting in accumulation into tissues and extracellular spaces is an important and regulated process of acute phosphate homeostasis, although the specific mechanisms involved were not resolved in this study. This finding is consistent with previous results demonstrating in healthy animals that urinary elimination of a phosphate challenge only achieved 50% by 4 hours, despite prior normalization of serum phosphate^14^. Similarly, in healthy humans, full clearance of a phosphate load required approximately 120 hours^15^ with no impact on circulating levels. Although the mechanisms responsible for prolonged, but not permanent phosphate storage, are not well understood, in CKD, this would likely have unique implications due to the likelihood of increased duration of storage when there is declining kidney function.

Adding tracer amounts of radiolabeled phosphate to a phosphate challenge facilitated the tracking of phosphate accrued in various compartments. In the CKD model, assessment of radioactivity in the chyme confirmed that active absorption was still occurring at 2 hours, but was nearly complete by 6 hours. Gut absorption of phosphate was not measurably altered by CKD or changes in dietary phosphate, which is similar to previous rat and human findings^8,16,17^. While it has been shown that intestinal NaPi2b is reduced by CKD and elevated dietary phosphate, functional assessments of phosphate absorption using whole tissue methods show no impairment in absorption indicating an importance of other compensatory mechanisms, such as Pit1/2 or paracellular transport in the setting of dietary phosphate sufficiency^18–20^.

In healthy animals the substantial deposition of new phosphate in the kidney cortex, liver, bone and cardiac muscle at 2 hours was further increased by 6 hours despite substantial urinary excretion during that time frame. In contrast, in healthy animals there was no further accrual into large arteries between 2 and 6 hours, suggesting a transient deposition in this tissue.

In both models of VC used in this study, significant blunting of the rise in circulating ^33^PO_4_ following the oral load was associated with substantial arterial accrual. In addition, increases in *de novo* phosphate accrual into blood vessels was associated with the amount of pre-existing VC. When the stimulus for calcification was removed in the calcitriol-mediated VC model, the phosphate accrual in the vessels was substantially attenuated and the circulating ^33^PO_4_, was similar to control animals. A strength of the study is that we were able to reproduce the findings in two different models of VC, where the stimulus and biomarker are markedly different, indicating that it is unlikely there is a direct role of uremic abnormalities or measured circulating factors associated with mineral metabolism. An exception is that FGF-23 that was elevated in both models, and was reduced when the stimulus for VC was removed in the calcitriol-mediated model. Both models are histologically similar and both have been reported to involve osteogenic transdifferentiation^21^: the upregulation of osteogenic markers (RUNX-2, osteocalcin) and loss of smooth muscle actin^22^, characteristics reflected in the human condition of medial calcification. This transition to a bone-like phenotype may explain the acquired active accrual capacity give calcified vasculature and bone had similar accrual per mg of tissue by 6 hours. Whether this transition and subsequent acute buffering of phosphate by the vasculature has physiological impacts on phosphate sensing by other organs (i.e. PTG or bone) or the normal circadian rhythm of phosphate is interesting and requires further study.

In the CKD animals, there was a substantial increase in bone deposition of *de novo* phosphate at 2 and 6 hours compared to control, which may be reflective of increased exposure of phosphate due to declined kidney function. Circulating PTH was much higher in the CKD animals on high phosphate, likely indicating increases in bone turnover, however this did not translate into impaired acute phosphate bone storage.

Intermittent exposure to phosphate may be an important stimulus of negative outcomes and VC progression, rather than hyperphosphatemia alone. First, it is recognized that medial vascular calcification occurs prior to chronic hyperphosphatemia in CKD patients^4^. Tani *et al*. show that chronic fluctuations in dietary phosphate induced much more VC than that same load of phosphate spread out consistently^6^, thereby indicating the importance of acute post-prandial elevations in serum phosphate The current understanding of medial calcification involves phosphate first entering the VMSCs through Pit-1, and then translocating to the extracellular matrix through pro-calcific Ca/Pi loaded vesicles^23^. Vascular Pit-1 is necessary for the development of VC and transdifferentiation of vascular smooth muscle cells (VSMCs) to an osteoblastic-like phenotype^24^. Transport-independent PiT-1 signalling can induce osteogenic differentiation, but the transport is required for extracellular matrix calcification^25^. This indicates that osteogenic differentiation is not sufficient for calcification, but needs to happen in conjunction with or secondary to, increased phosphate influx, potentially of the type depicted in this study. While Pit-1 expression has been shown to increase in response to elevated dietary phosphate in CKD^26^, upregulation of PiT-1 alone is not sufficient for VC, and must happen in conjunction with increased circulating phosphate, as demonstrated by a transgenic mouse model with upregulated PiT-1, where VC was not present^27^. These fluctuations in phosphate also have negative consequences on endothelial cells, increasing oxidative stress and inflammatory responses^28^, both of which have negative consequences on cardiovascular health^3^. Phosphate fluctuation are a potential mechanism through which more frequent and longer dialysis confer positive health outcomes, although this has not been well-examined as through medial VC as a mediator^29^.

CKD animals fed a low phosphate diet and lacking measurable calcification had profiles of vascular deposition at both the 2- and 6-hour time points that were similar to controls despite the acute change in circulating phosphate similar to CKD high phosphate. In other words, the circulating phosphate stimulus for deposition is similar but accrual or retention was impaired. This finding is unlikely to be due to impaired cellular transport, as there is no evidence suggesting a downregulation of Pit-1 in CKD, and even uremic toxins alone have been shown to upregulate Pit-1^30^. It could however be a result of impaired retention of phosphate through upregulated XPR1, the major phosphate export protein, whereby dysfunction leads to brain calcifications^31^, although it’s role in VC had not been studied. Alternatively, this finding may indicate a successful role calcification inhibitors in making the microenvironment less favourable to phosphate deposition, such as fetuin A, pyrophosphate, or matrix gla protein^32–34^.

Comparison of the phosphate challenge to a radiolabelled saline equivalent allowed us to compare active acute accrual to passive equilibration with the exchangeable pool of phosphate in the tissues. Compared to the saline load, which contained similar tracer levels of ^33^PO_4_, the phosphate challenge produced substantially more accrual into the calcified vessels, but not in the non-calcified vessels. The differential acute accrual of phosphate into only the blood vessels supports the concept that once VC is initiated, it progresses quickly and potentially through a different mechanism than initiation. In incident dialysis patients, only patients with pre-existing calcification had significant progression over the first 18 months^35^.

These findings have important consequences for dietary management of CKD patients. Based on 2001-2014 NHANES data, the average adult consumes at least twice the recommended daily intake of phosphate^36^, whereby approximately 50% is estimated to be derived from high bioavailable inorganic food additives^37,38^. Inorganic sources, such as food additives, which are 90-100% absorbed, as compared to plant- or meat-derived phosphate, which is much lower (40-69%)^39^. These inorganic sources of phosphate are quickly absorbed, leading to a more rapid flux into the circulating pool. Patients with CKD and hyperphosphatemia are instructed to consume low phosphate diets and prescribed intestinal phosphate binders. Phosphate binder therapy slows the progression of VC, but some sub analyses have shown its most dramatic effect is on the progression of pre-existing calcification^40^.

In the control HP animals there was a very significant correlation of *de novo* deposition in the vasculature at 6 hours compared to resident phosphate, despite the lack of VC, that wasn’t present in the LP controls. This finding potentially indicates that dietary phosphate, even in the setting of healthy kidney function, alters the vascular handling of phosphate, a finding that would need to be confirmed in larger studies.

The novel whole-body physiological approach to assessing the acute phosphate response allowed comprehensive assessment of tissue deposition, and represents a powerful tool to assess the sequential steps leading to VC and the impact of vascular-specific interventions. There is limited evidence to suggest that calcification can regress, so nuanced assessments of activity at different stages will be important for assessing treatments aimed at limiting progression. The current study was limited in that 1) we did not elucidate the specific mechanism by which this acute accrual occurs and 2) whether the accrual in this study represents long-term deposition or temporary storage (i.e. how much, if any, of the acute vascular deposition translated into increased accrual of VC), which are important area of future research.

In summary, this study characterized the role of calcifying arteries in the acute non-renal clearance of phosphate following a phosphate load in two experimental models of VC. Our data indicate that calcifying arteries alter the systemic disposition of a phosphate challenge and acutely deposit substantial phosphate. This study supports the importance of diet as it relates to acute fluctuations of circulating phosphate and the importance of bioavailability and meal-to-meal management in CKD patients as a mediator of cardiovascular risk.

## Acknowledgements

Study Design: MET, JGEZ, BAS, MAA, RMH. Study Conduct: MET, APL, PSJ, LHL. Data Collection: MET, APL, PSJ, LHL. Data Analysis: MET, MAA, RMH. Data Interpretation: MET, MAA, RMH. Drafting Manuscript: MET, APL, MAA. Revising Manuscript and Content: MET, APL, MAA, RMH. Approving final version of manuscript: MET, APL, PSJ, LHL, BAS, JGEZ, RMH, MAA. MET takes responsibility for the integrity of the data analysis.

