## Supplemental Tables and Figures for "Vascular calcification has a role in acute non-renal phosphate clearance" (from another version of this preprint: version 1, 2020-07-29)

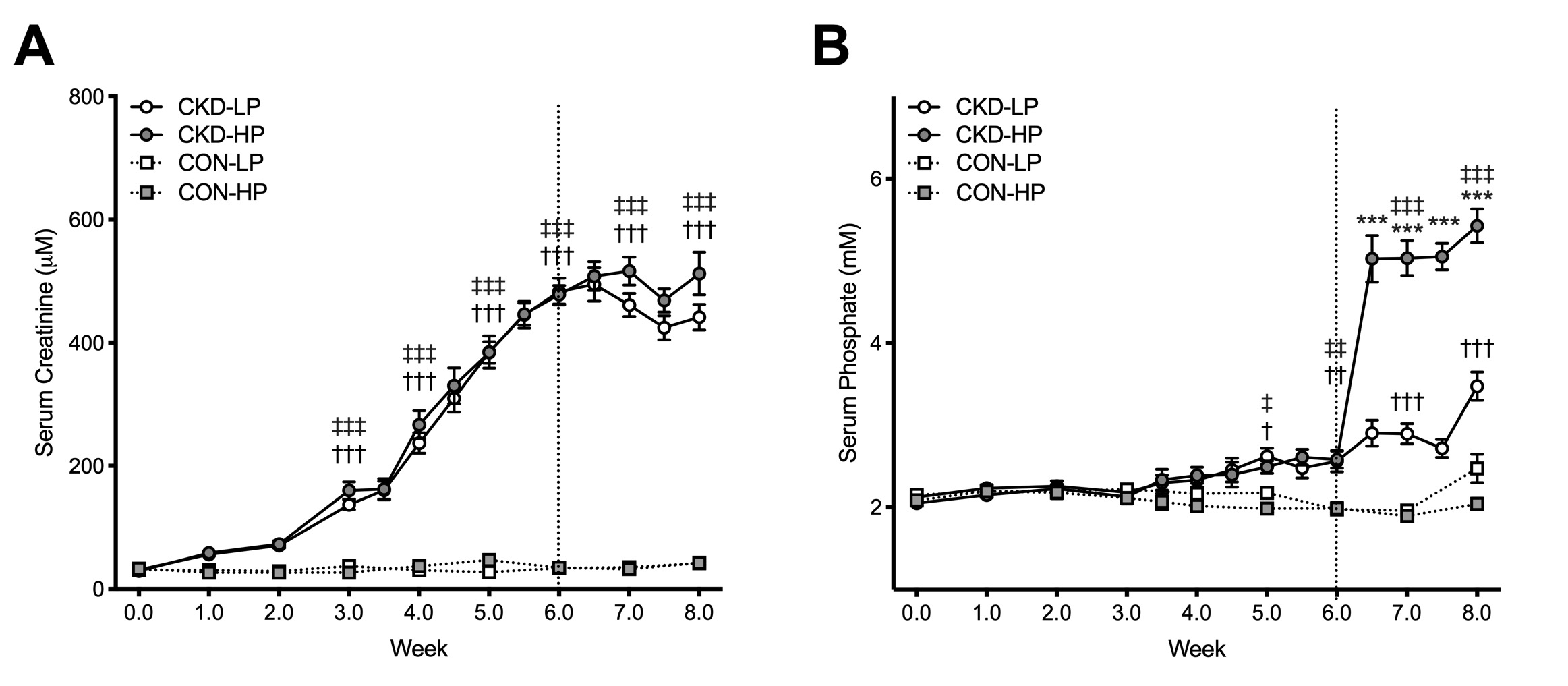
**Supplementary Figure 1: Increased circulating creatinine (A) and phosphate (B) during the development of adenine-induced CKD.** Increased severity of hyperphosphatemia but not creatinine levels with exposure to high dietary phosphate. Differences assessed by two-way ANOVA and *post hoc* Tukey corrected T-tests. Double daggers indicate differences between CKD-HP and CON-HP, single daggers indicated a difference between CKD-LP and CON-LP. * indicate a difference between CKD-LP and CKP-HP. *p<0.05, **p<0.01, ***p<0.001

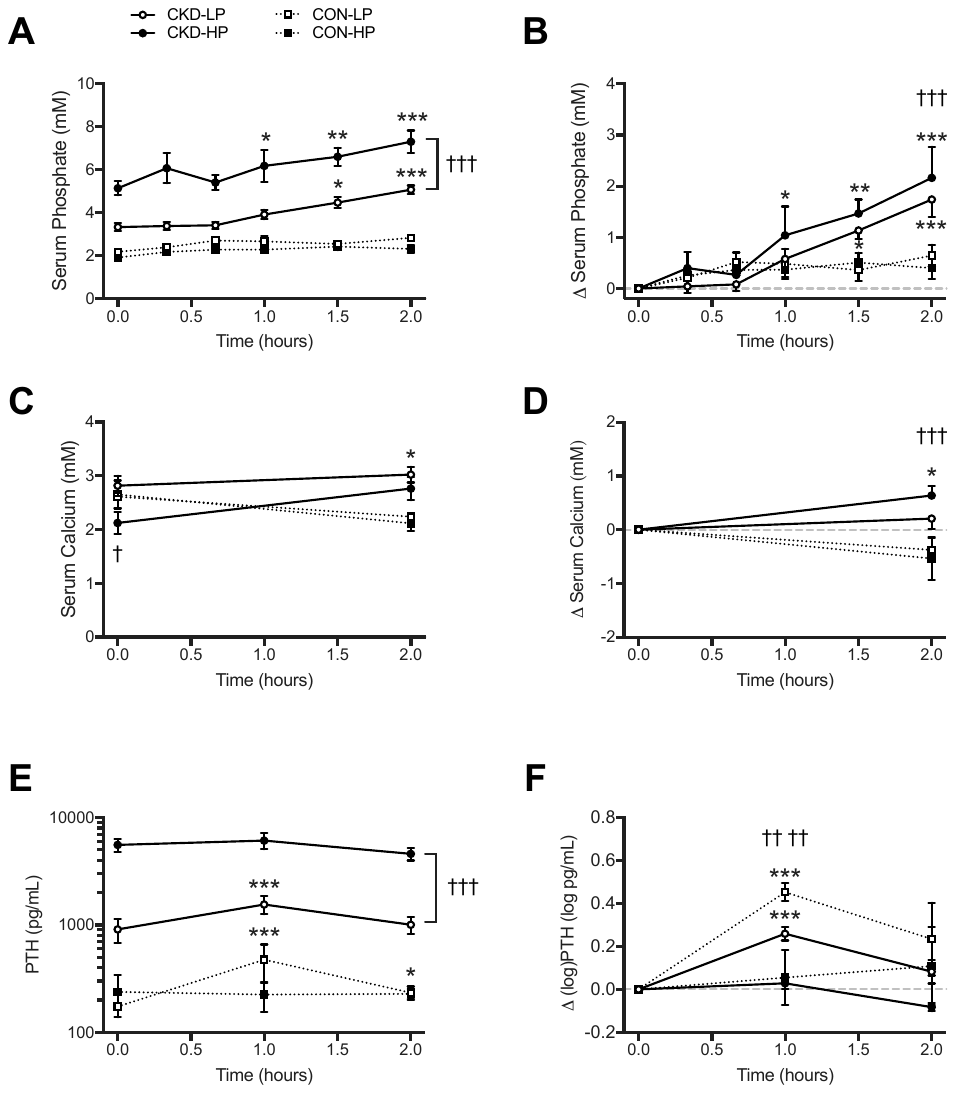

**Supplementary Figure 2:** Circulating measures following oral phosphate load in animals sacrificed at 2 hours. Phosphate (A), absolute change in phosphate (B), calcium (C), absolute change in calcium (D), PTH (E) and absolute change in log-transformed PTH (F). Repeated measures two-way ANOVA evaluating within-group differences from time 0 (Dunnett’s correction; * p<0.05, *p<0.01, ***p<0.001) and between group differences comparing CKD-LP to CKD-HP (Tukey correction; † p<0.05, ††† p<0.001). Data expressed mean ± SEM. Comparisons in PTH evaluated on log-transformed data.

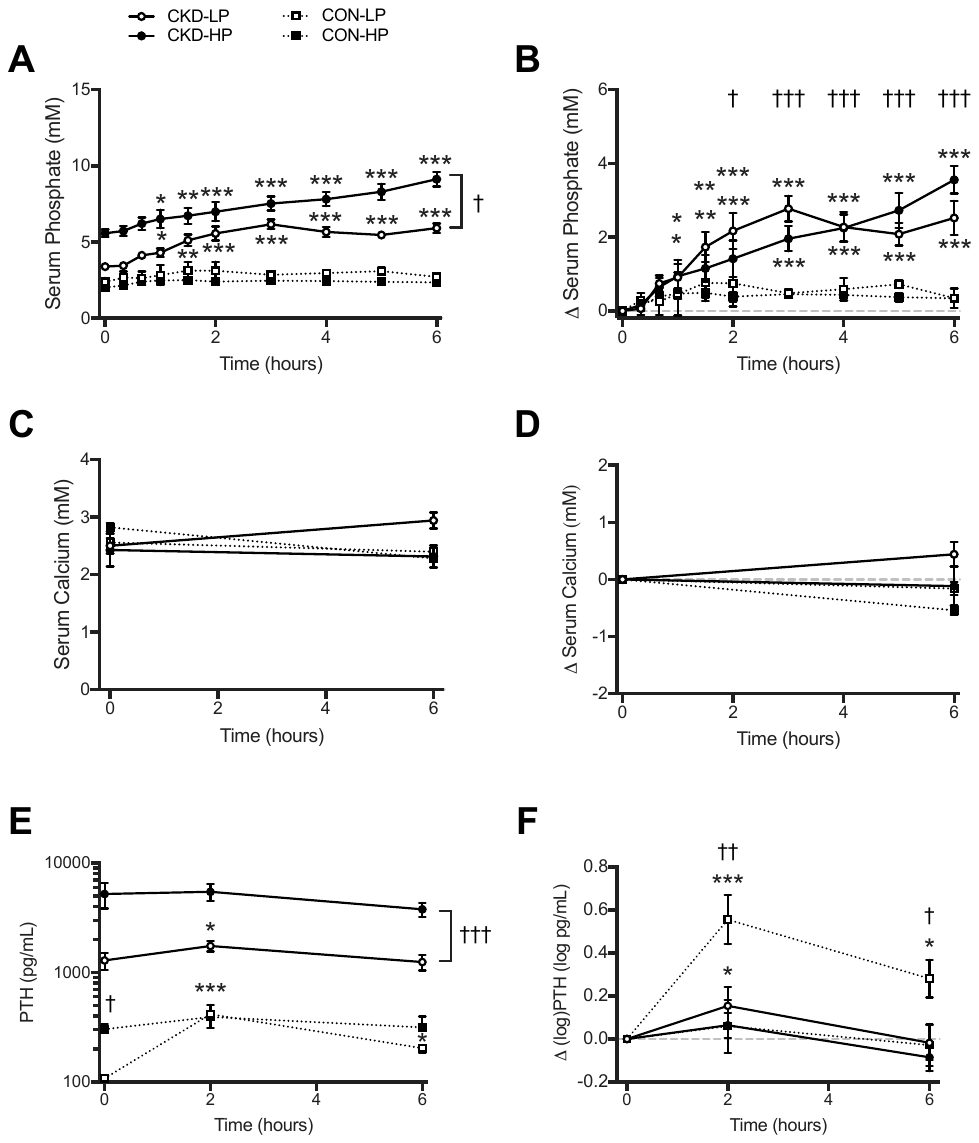

**Supplementary Figure 3:** Circulating measures following oral phosphate load in animals sacrificed at 6 hours. Phosphate (A), absolute change in phosphate (B), calcium (C), absolute change in calcium (D), PTH (E) and absolute change in log-transformed PTH (F). Repeated measures two-way ANOVA evaluating within-group differences from time 0 (Dunnett’s correction; * p<0.05, *p<0.01, ***p<0.001) and between group differences comparing CKD-LP to CKD-HP (Tukey correction; † p<0.05, ††† p<0.001). Data expressed mean ± SEM. Comparisons in PTH evaluated on log-transformed data.

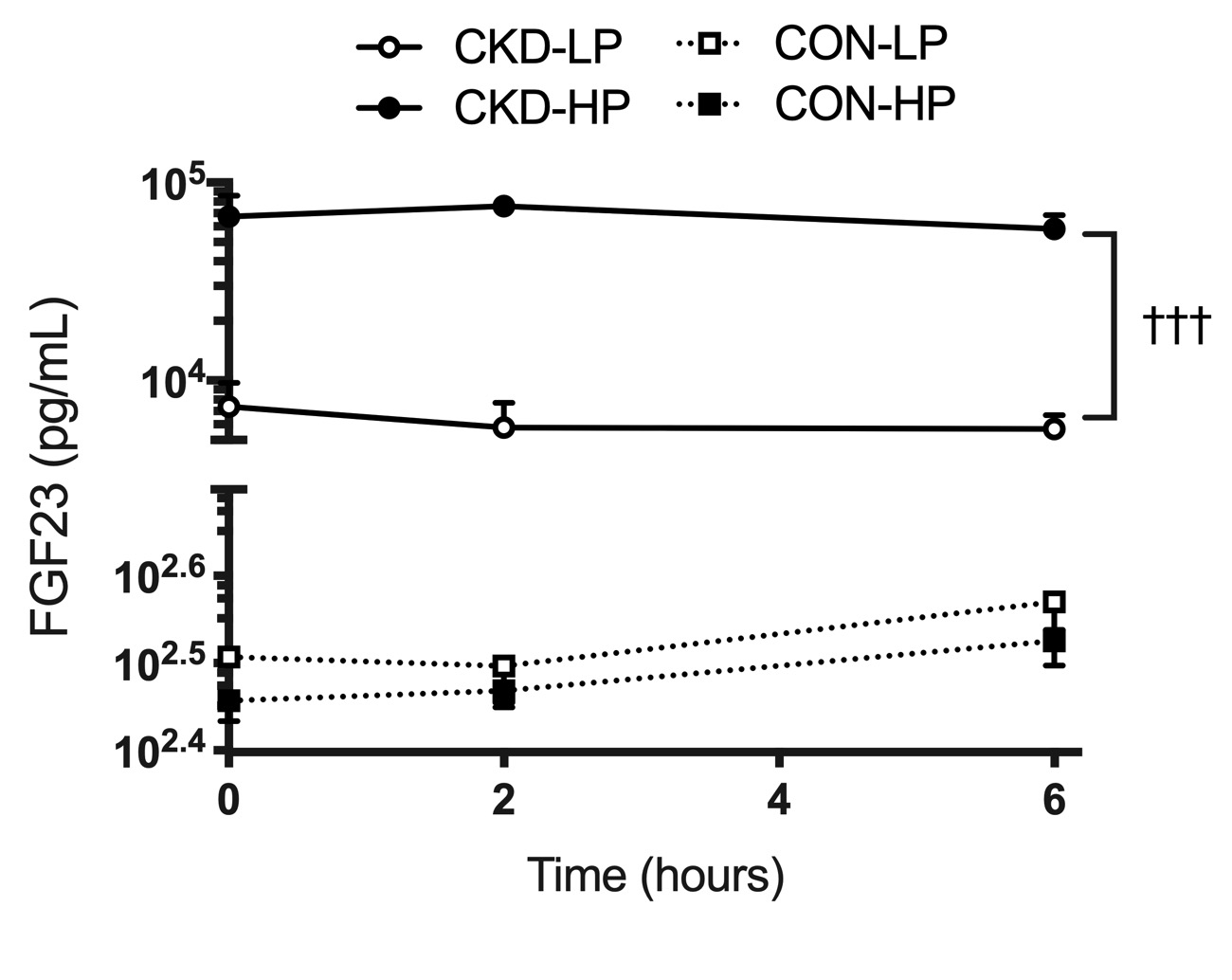

**Supplementary Figure 4: Minimal change in circulating of FGF-23 two and 6 hours after at sacrifice in animals given the phosphate.** Two-way ANOVA evaluating within-group differences from time 0 (Dunnett’s correction; * p<0.05, *p<0.01, ***p<0.001) and between group differences comparing CKD-LP to CKD-HP (Tukey correction;† p<0.05, ††† p<0.001). Data expressed mean ± SD. Comparisons evaluated on log-transformed data.

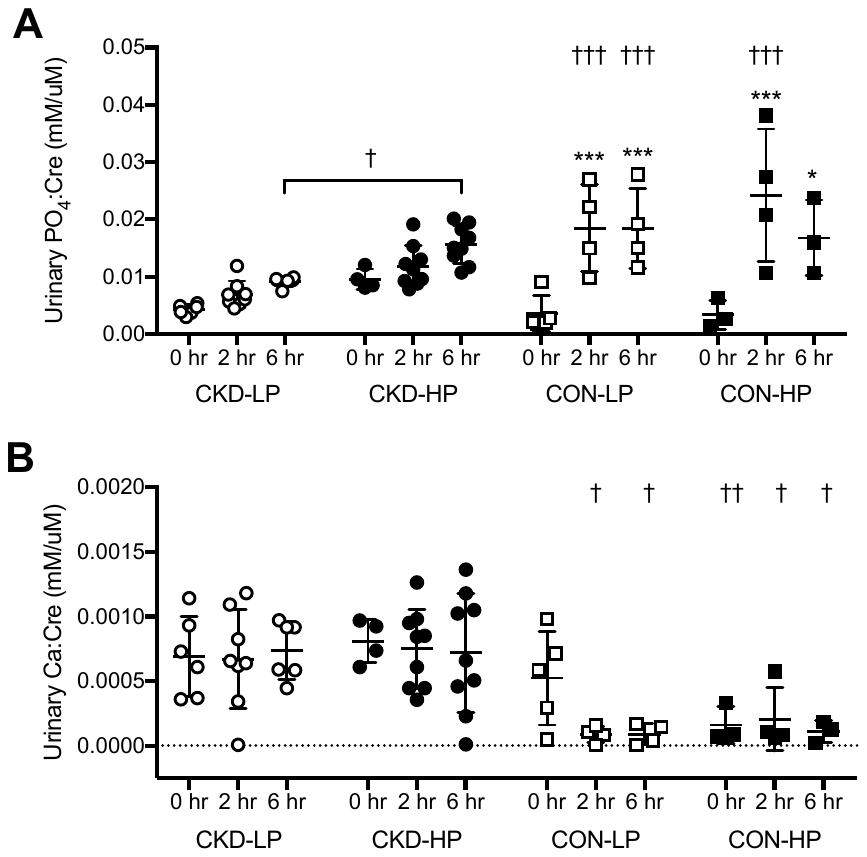

**Supplementary Figure 5:** Urinary calcium to creatinine ratios at sacrifice. Two-way ANOVA evaluating within-group differences from time 0 hr (Dunnett’s correction; * p<0.05, ***p<0.001) and between group differences at each time point (Tukey correction; † p<0.05, ††† p<0.001). Unless otherwise indicated, symbols reflect changes between diet-matched CON v CKD. Data expressed mean ± SD.

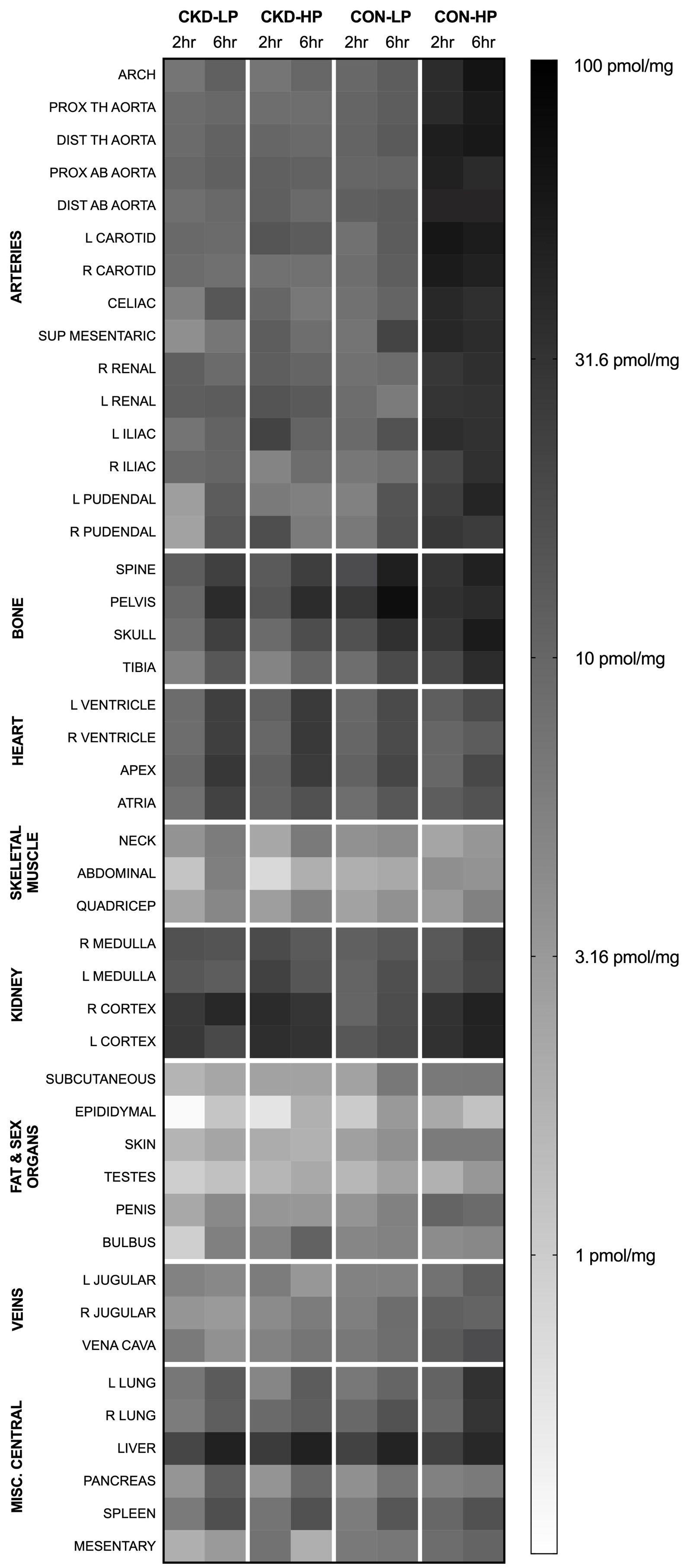

**Supplementary Figure 7:** **Experimental CKD and dietary phosphate alter tissue disposition of an oral load of radiolabeled phosphate.** *De novo* tissue phosphate accrual in various tissues across the body at 2 and 6 hours following oral load grouped by tissue type. The heat map coloration represents the amount of *de novo* phosphate accumulated per mg of wet-weight tissue on a logarithmically-transformed scale. Arteries sorted by external diameter. For full tissue list see supplementary methods.

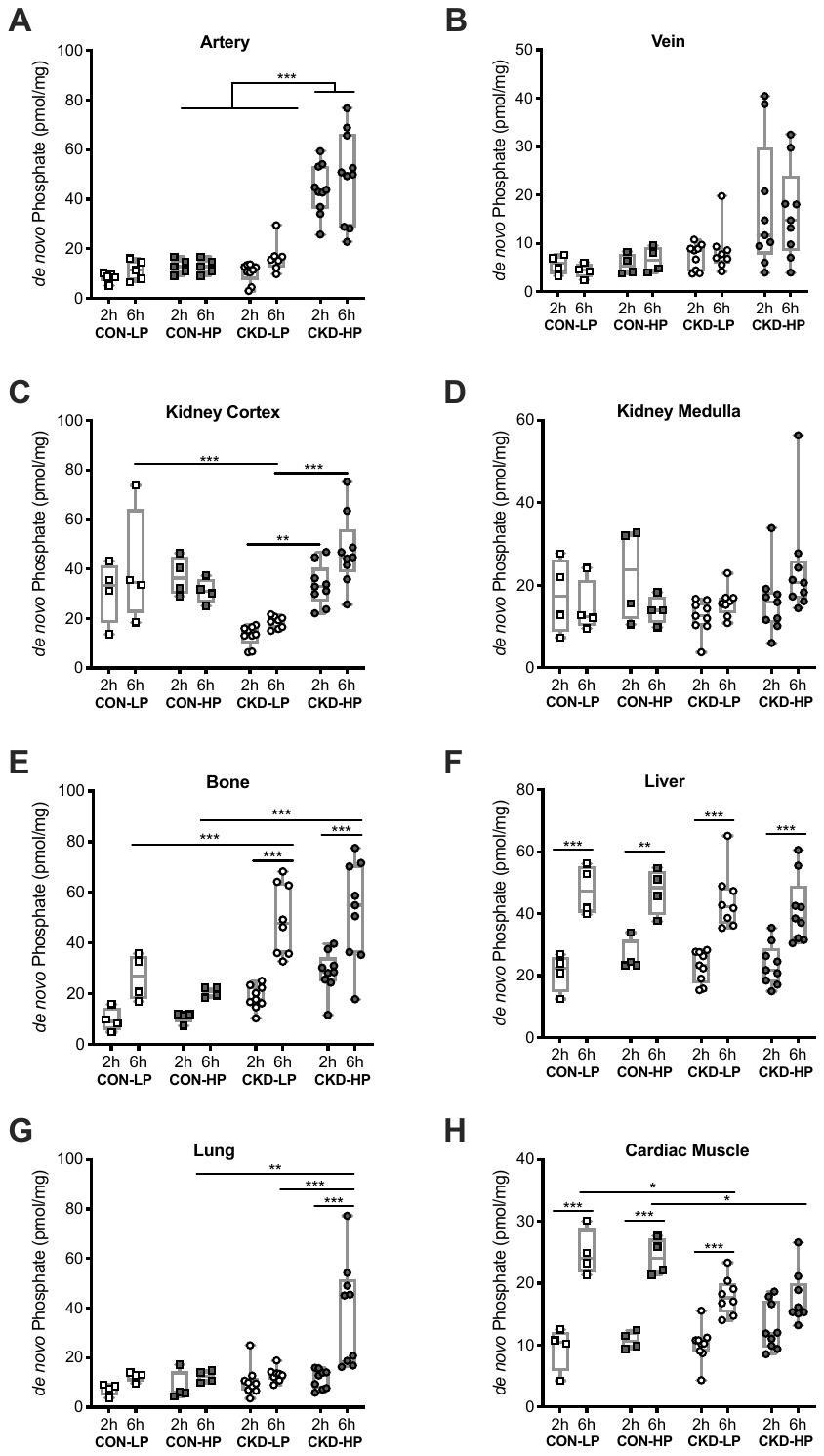

**Supplementary Figure 8:** *De* novo tissue phosphate accrual in **(A)** arteries, (B) veins, (C) kidney cortex, (D) kidney medulla, (E) bone, (F) liver, (G) lung, and (H) cardiac muscle. Each data point represents the pooled average of samples for each tissue type for each individual rat, as indicated in Figure 2H. Three-way ANOVA with *post hoc* Sidak-corrected multiple comparisons. *p<0.05, **p<0.01 *** p<0.001 between data-sets that differed only by one variable.

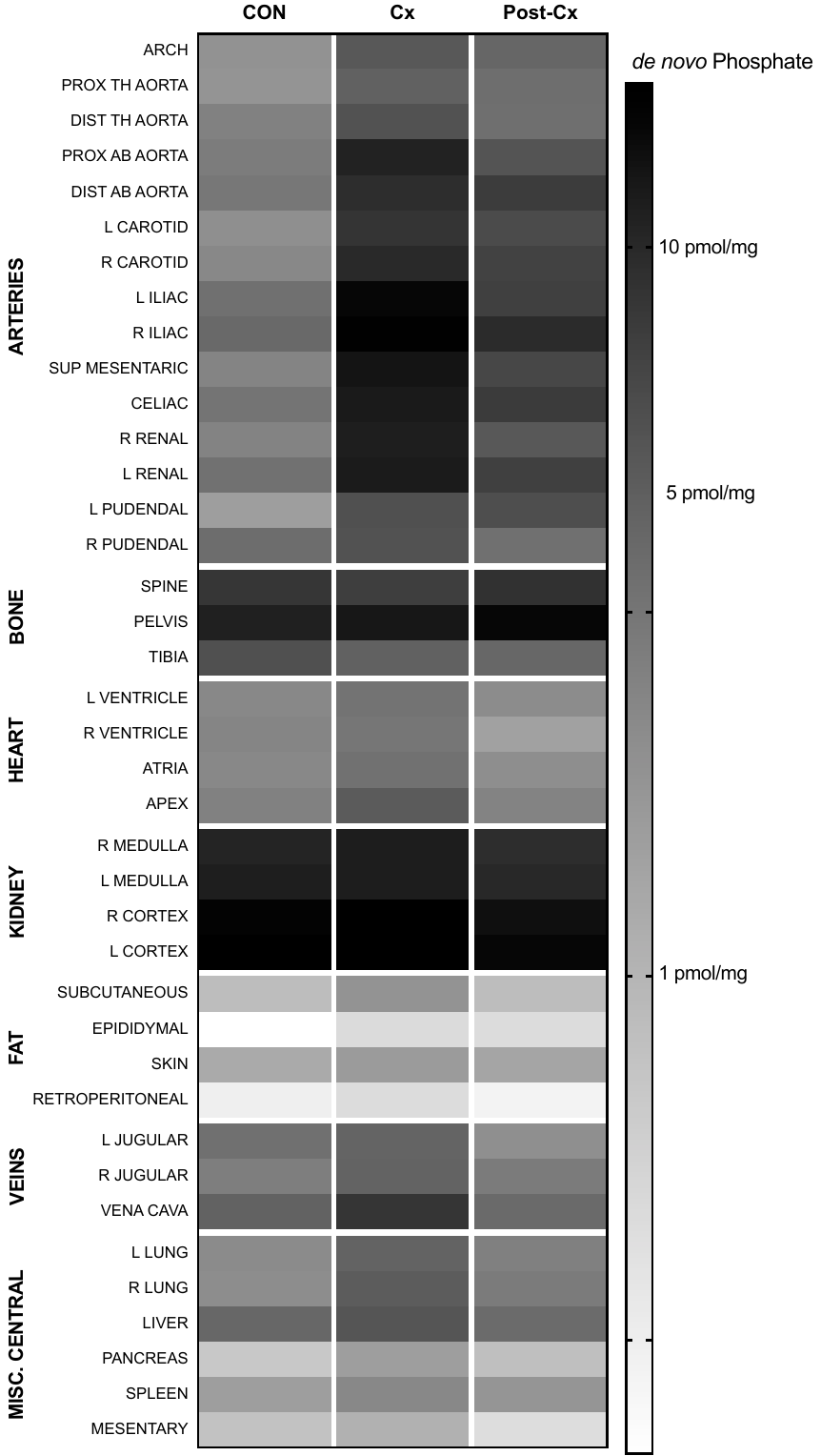

**Supplementary Figure 9:** **Calcifying vasculature alters tissue disposition of an IV radiolabeled phosphate challenge.** *De novo* tissue phosphate accrual in various tissues across the body at 2 and 6 hours following oral load grouped by tissue type. The heat map coloration represents the amount of *de novo* phosphate accumulated per mg of wet-weight tissue on a logarithmically-transformed scale. Arteries sorted by external diameter. For full tissue list see supplementary methods.

Supplementary Table 1: Model Characteristics at Sacrifice between treatment groups and time-points of sacrifice

| **Treatment Group** | **Time of Sacrifice** | **SAC Creatinine**  **(μM)** | **7.5 Wk Creatinine**  **(μM)** | **Baseline PO_4_**  **(mM)** | **Baseline Ca**  **(mM)** | **Baseline PTH (pg/mL)** | **Baseline PTH**  **(pg/mL) Med[IQR]** | **(LOG) Baseline PTH (log pg/mL)** | **Bodyweight**  **(g)** |
| --- | --- | --- | --- | --- | --- | --- | --- | --- | --- |
| **CKD 0.5%** | **ALL** | **441.4±100.0 ^4^*,^4^^,** | **424.3±94.0 ^4^*,^4^^,** | **3.74±0.80 ^3^*,^4^^** | **2.79±0.52** | **1039±651** | **930 [542,1479] ^2^*,^4^^** | **2.94±0.28 ^4^*,^4^^** | **465±27 ^3^*,^,** |
|  | *0 hr (N=6)* | 416.5±71.8 | 428±71.8 | 3.77±1.33 | 3.13±0.53 | 899±620 | 990 [378,1516] | 2.89±0.31 | 467±27 |
|  | *2 hr (N=9)* | 464.7±142.7 | 431.0±117.1 | 3.33±0.50 | 2.82±0.53 | 909±687 | 759 [369,1258] | 2.86±0.21 | 476±26 |
|  | *6 hr (N=8)* | 434.0±56. | 413.2±78.9 | 3.39±0.57 | 2.50±0.39 | 1290±643 | 1333 [690,1503] | 3.06±0.03 | 459±27 |
| **CKD 1.0%** | **ALL** | **512.4±166.6 ^4^*,^4^^,** | **468.7±89.5 ^4^*,^4^^,** | **5.05±0.98 ^4^*,^4^^,^4^**^δ^ | **2.47±0.72** | **4688±3143 ^4^*,^4^^,^4^**^δ^ | **3100 [2497, 6090] ^4^*,^4^^,^2δ^** | **3.95±0.25 ^4^*,^4^^,^4^**^δ^ | **411±25 ^4^*,^4^^,^4^**^δ^ |
|  | *0 hr (N=5)* | 531.1±202.2 | 468.6±88.0 | 5.68±1.33 | 3.08±0.56 | 2342±186 | 2339 [2163, 2523] | 3.37±0.17 | 401±25 |
|  | *2 hr (N=9)* | 466.1±88.4 | 452.2±52.4 | 5.14±1.00 | 2.13±0.57** | 5559±2178** | 5324 [3885, 7458]** | 3.71±0.30 (p=0.07) | 408±20 |
|  | *6 hr (N=9)* | 510.9±189.3 | 485.±121.4 | 5.58±0.77 | 2.43±0.76 (p=0.06) | 5216±4154* | 3063 [2656, 8285] | 3.62±0.46 | 417±30 |
| **CON 0.5%** | **ALL** | **41.7±8.8** | **41.7±8.8** | **2.47±0.62** | **2.51±0.28** | **252±379** | **131 [99, 220]** | **2.19±0.37** | **506±22** |
|  | *0 hr (N=5)* | 37.2±5.1 | 37.2±5.1 | 2.78±0.81 | 2.39±0.07 | 429±588 | 191 [114, 863] | 2.39±0.46 | 503±24 |
|  | *2 hr (N=4)* | 41.6±6.1 | 41.6±6.1 | 2.18±0.27 | 2.61±0.43 | 179±158 | 117 [67, 338] | 2.12±0.36 | 513±21 |
|  | *6 hr (N=4)* | 47.4±12.6 | 47.4±12.6 | 2.37±0.55 | 2.56±0.29 | 108±26 | 111 [81. 131] | 2.02±0.11 | 504±25 |
| **CON 1.0%** | **ALL** | **42.8±10.4** | **42.8±10.4** | **2.04±0.23** | **2.57±0.31** | **241±131** | **243 [109, 296]** | **2.31±0.27** | **497±28** |
|  | *0 hr (N=3)* | 30.7±6.5 | 30.7±6.5 | 2.25±0.31 | 2.36±0.12 | 162±47 | 181 [109, 131] | 2.19±0.14 | 502±32 |
|  | *2 hr (N=4)* | 41.7±6.7 | 41.7±6.7 | 1.91±0.08 | 2.60±0.38 | 238±71 | 194 [73, 197] | 2.24±0.41 | 499±35 |
|  | *6 hr (N=4)* | 52.9±1.2 | 52.9±1.2 | 2.02±0.19 | 2.86±0.16 | 303±66 | 287 [251, 448] | 2.27±0.09 | 492±23 |
| **ANOVA: Treatment Group** | | **p<0.0001** | **p<0.0001** | **p<0.0001** | **p=0.2** | **p<0.0001** | **p<0.0001** | **p<0.0001** | **p<0.0001** |
| **ANOVA: Time of Sacrifice** | | **p=0.6** | **p=1** | **p=1** | **p=0.4** | **p=0.4** | **p=0.4, p=0.06, p=0.4, p=0.3** | **p=0.6** | **p=0.70** |

Differences between treatment group means: Tukey’s multiple comparisons test between each treatment group. * Difference from CON 0.5% ^ Difference from CON 1.0%

^δ^ Difference from CKD 0.5%

Control 7.5 week creatinine is actually 8 week as they were only sampled weekly for measurements.

PTH: CKD 1.0% and CON 0.5% = not normally distributed (D’Agostino)

PTH med[IQR] Comparison between treatment groups: Kruskal Wallis, Dunn’s Multiple comparisons (same thing for between each time of sacrifice within each treatment group)

FGF-23 values are taken at sacrifice after the phosphate infusions, not prior to the phosphate infusion.
