## Supplemental Methods for "Vascular calcification has a role in acute non-renal phosphate clearance" (from another version of this preprint: version 1, 2020-07-29)

**Complete Methods**

***Adenine-Induced CKD Model and Oral Radioactive Phosphate:***

*Adenine-Induced CKD Model:* All animal procedures were performed in accordance with the Canadian Council on Animal Care and were approved by Queen’s Animal Care Committee. Male Sprague-Dawley rats (N=67: Hilltop Lab Animals Inc. PA, USA),15-16 weeks of age, were acclimated for a week prior to experimental start and were individually housed and maintained on a 12-hour light/dark cycle throughout the duration of the study. Progressive kidney dysfunction was induced using a dietary adenine model, as previously described with modifications . In brief, animals were stratified at study start by weight into either a control(CON) or chronic kidney disease group using specially formulated diets that were identical, aside from the addition of 0.25% adenine in the CKD group. The diet contained 0.5% phosphate, 1% calcium, 0.05% magnesium, 0.2 mg/kg vitamin K, 1 IU/g cholecalciferol, and 6% protein (Harlan Tekland, WI, USA: 0.25% Adenine, 0.5% P: TD.08672, 0.5% P: TD.150555, 1.0% P: TD.08670). Adenine diet and water was fed *ad libitum.* Control diet was provided to a maximum of 20g/day. Animals were maintained on this adenine supplemented diet for 4.5-5.5 weeks, depending on the extent of creatinine elevation. Animals were sampled 2x weekly and removed from adenine after two consecutive measures of serum creatinine >300uM. After removal of the adenine diet, animals were maintained on the non-adenine 0.5% phosphate diet for at least 4 days to ensure removal of the acute effects of dietary adenine. At the sixth week, CKD and CON rats were stratified into high or low dietary phosphate. Stratification was based on serum creatinine, serum phosphate, calcium and bodyweight. The low dietary phosphate group remained on the 0.5% dietary formulation (LP) and the high dietary phosphate (HP) increased to 1% dietary phosphate (TD.08670). At this point, all animals begun scheduled feeding at 10g twice per day until sacrifice. When animals did not eat all given food or reached 10% weight loss, they were supplemented with a high calorie supplement (Boost®, Clear H2O, USA) supplemented with mono-basic sodium phosphate to ensure consistent dietary phosphate consumption. The total number of animals in each group are: CON-LP (N=13), CON-HP (N=11) CKD-LP (N=23) and CKD-HP (N=23). The discrepancy in control rat numbers was due to a labelling error.

*Oral Phosphate Sacrifice Protocol:* At 8 weeks, animals were prepared for sacrifice. Animals were given 15g of food the day before sacrifice to provide a partially-fasted state to ensure consistency of stomach contents during the morning protocol described below. At approximately 7-10am, animals were provided 2mL of sucralose gel (MediGel®, Clear H2O, USA) with a total phosphate amount of 0.1g (equivalent to 100% daily intake of the LP animals, or 50% daily intake of HP animals). Phosphate in the gel was supplemented with dibasic and monobasic sodium phosphate salts (Sigma-Aldrich, Canada). The sucralose was also spiked with ~7.79 million Bq of radio-labeled ^33^PO_4_ (NEN Radiochemicals, Perkin Elmer, USA). Animals were stratified by three most recent measurements of serum creatinine, phosphate, calcium and bodyweight into one of three sacrifice times following the oral load of phosphate: 0 hour, 2 hours or 6 hours. Stratification metrics and final study animal numbers for each time point is outlined in Supplemental Table 1. Depending on sacrifice time, animals were blood sampled at the saphenous vein at times 0, 20min, 40min, 1hr, 1.5hr and 2hr and then hourly until 6hr.

***Calcitriol-Induced Model of Vascular Calcification and Radioactive IV Phosphate Challenge:***

*Calcitriol Daily Injection Model of Vascular Calcification:* All animal procedures were performed in accordance with the Canadian Council on Animal Care and were approved by Queen’s Animal Care Committee. Male Sprague-Dawley rats (N=24: Hilltop Lab Animals Inc. PA, USA),14 weeks of age, were acclimated for a week prior to experimental start and were individually housed and maintained on a 12-hour light/dark cycle throughout the duration of the study. Two-days before interventions, animals were transferred from standard rat chow (Purina) to a specially formulated diet that contained 0.75% phosphate, 1% calcium, 0.05% magnesium, 0.2 mg/kg vitamin K, 1 IU/g cholecalciferol, and 6% protein (Harlan Tekland, WI, USA: TD.160324). Water was fed *ad libitum.* Animals were stratified at study start by weight into either a control or treatment groups. Treatment rats were injected subcutaneously with calcitriol (0.5 ug/kg bodyweight) once per day (~8am) for 8 days (Sterimax, ON, CAN). Throughout the experiment, control dietary intake was matched with treatment rat intake to maintain bodyweight and nutrient intake similarity. When animals did not eat all given food or reached 10% weight loss, they were supplemented with a high-calorie supplement (Boost®, Clear H2O, USA).

Animals were blood sampled from the saphenous vein at 8am before dosing every other day beginning at baseline, prior to the first dose. Sampling was done with a 22G needle and microcapillary blood collection tubes (2 whole blood, 4 heparinized plasma). Whole blood was left at room temperature for at least 45 mins prior to centrifugation to generate serum. Heparinized whole blood was collected on ice prior to centrifugation. Samples were centrifuged at 9,000 RPM for 20 minutes at 4°C. Calcium, phosphate, PTH and FGF-23 were measured at each sample collection.

Following the blood sample on day 7 of treatment, both treatment and control animals were stratified into two groups based on serum calcium, phosphate, PTH, FGF-23, and bodyweight. The first group was? sacrificed on experimental day 9, following 8 doses of calcitriol (Cx). The remaining rats no longer received calcitriol (and controls) for 13 days (Post-Cx). The total number of animals in each group are: Cx (N=8), Post-Cx (N=9), Control Early (N=3), and? Control Late (N=4).

In brief, animals were administered an intravenous infusion of a phosphate load spiked with radiolabeled phosphate to track tissue-based deposition and movement. Intravenous delivery was chosen over oral administration as the delivery method due to the impact of calcitriol on intestinal phosphate absorption. Animals were anesthetized with inhaled isoflurane (5% isoflurane, 2% O_2_) in an induction chamber and brought to surgical plane with reduced isoflurane (2.5-3%) via nose-cone. A baseline blood sample was taken from the saphenous vein. A neck incision exposed the right and left jugular vein. A venous infusion PE-50 cannula was inserted into the left jugular (right jugular as a second option).

Sodium mono-basic and di-basic phosphate (Sigma-Aldrich, Canada) was dissolved to generate a solution with a pH of 7.4. Each rat was infused with a total of 3mL of solution totaling 300μmol of phosphate, and ~9.7 million Bq/mL, NEN Radiochemicals, Perkin Elmer, USA). Infusate was warmed to 37°C and infused using an infusion pump (KD Scientific) at a rate of 0.3mL/min for 10 minutes. Blood was sampled directly post-infusion at 10 minutes (from the jugular vein), 20 minutes and sacrifice (30 minutes).

A subsequent study was completed administering calcitriol subcutaneously via osmotic minipump. Aside from method of administration, all other methodologies were identical to previous study. Male Sprague-Dawley rats (N=12: Hilltop Lab Animals Inc. PA, USA),14 weeks of age underwent surgical implantation of subcutaneous Alzet osmotic minipumps (2mL capacity, 10μL/hr flow rate) filled with calcitriol solution (Sterimax, ON, CAN). The calcitriol solution was diluted with saline to concentration based on the flow rate of the pump and the animal’s preoperative bodyweight, corresponding to a cumulative daily dose of 0.5μg/kg bodyweight. Prior to implantation, pumps were primed in saline at 37°C. Animals were anesthetized with inhaled isoflurane (5% isoflurane, 2% O_2_) in an induction chamber and brought to surgical plane with reduced isoflurane (2.5-3%) via nose-cone. A two-inch incision was made along the animals back and a small subcutaneous pocket was made using blunt dissection. Following insertion of the pump into the pocket, the incision was sutured. Animals were given a postoperative loading dose of subcutaneous meloxicam (2mg/kg bodyweight) in 10mL saline followed by 0.6mL of lidocaine:bupivicane given intra-dermally along the incision. 24 hours after the initial meloxicam injection, animals were given postoperative subcutaneous meloxicam (1mg/kg bodyweight) in 10mL saline. This scheduled postoperative treatment was continued for 3 days following surgery. Following the blood sample on day 7 of treatment, animals were stratified into two groups based on serum calcium, phosphate, PTH, FGF-23 and bodyweight. The first group (N=6) received the intravenous infusion of a phosphate load spiked with radiolabeled phosphate as described above. The second group (N=6) received an infusion of only the radiolabeled phosphate, made up in saline and lacking the 300μm phosphate load.

***Sacrifice Protocol and Radioactivity Assessment***

Animals were anesthetized with isoflurane (5%) and sacrificed via cardiac puncture and exsanguination. Urine was collected directly from the bladder. Gastrointestinal tissue from the stomach to anus was quickly excised and separated. Samples of chyme were collected from the stomach, proximal small intestine (duodenum) and distal small intestine (ileum) and large intestine. Feces was collected from the distal colon. Various somatic tissues were collected: (50) including various samples of arteries, veins, cardiac and skeletal muscles, bone, kidney, fat, intestine, liver, pancreas, lung. The full list of tissues collected is appended below. Tissues were rinsed in saline and patted dry to remove any blood prior to measurement. Tissues and chyme was put into 1N HCl at a ratio of 50uL/mg for one week at room temperature. After one week, samples were vortexed and centrifuged for 10 minutes. The acidic, clear top layer was separated from the pellet.

*Biochemical blood and urinary measurements:* Serum creatinine and serum and urine calcium and phosphate were evaluated spectrophotometrically (SynergyHT Microplate Reader; BioTek Instruments, Winooski, VT). Creatinine was evaluated using the Jaffe method (QuantiChrom^TM^ Creatinine Assay Kit, Bioassay Systems). Calcium was measured using the o-cresolphthalein complexone method at 540nm, whereby the calcium in the sample forms a purple complex with the O-cresolphthalein colour reagent (Sigma-Aldrich, CAN)^2^. Free phosphate was measured using the malachite green (Sigma-Aldrich, CAN) method as described by Heresztyn and Nicholson^3^ at 650nm. Plasma levels of intact PTH and C-terminal /IntactFGF-23 were measured using ELISAs (Immunotopics Inc., USA). Tissue levels of calcium and phosphate were measured per mg of wet weight using the same assays above with pH-controlled standards.

*Radioactivity measurement and analysis:* A week after sacrifice, ^33^PO_4_ counts per minute (CPM) were measured in tissue and chyme acid homogenate (obtained as outlined above), serum, urine as well as an aliquot of the solution given to each rat. Adding an aliquot of the solution given to each rat with the measurements of serum and tissue allowed us to account for radioactive decay occurring throughout the study. Samples (100-200uL) were added to ~5 mL Ultima Gold AB scintillation cocktail (Perkin Elmer, USA) and analyzed using a Beckman Coulter LS 6500 multi-purpose scintillation counter. Each sample was measured twice for a 1-minute count time. Corrected CPM were obtained by subtracting background from the CPM for all samples and then normalized to each rat’s ingested radioactivity.

In order to transform counts/mg of tissue to an estimation of amount of phosphate accrued per tissue, the following analysis was done. Serum specific activity was calculated at each time point over the course of the study (equation 1). A time-weighted average serum specific activity was generated and was used to estimate tissue phosphate accrual (equation 2). CPM were transformed into Bq using the ^33^P efficiency of the scintillation counter.

(1) Serum Specific Activity (CPM/pmol) = Serum Radioactivity (CPM/uL) / Serum Phosphate (mM)
(2) Tissue PO_4_ Accrual (pmol PO4/mg tissue) = Tissue Radioactivity (CPM/mg tissue) / Average Specific Activity (CPM/pmol)

*Von Kossa Method of Visualizing Vascular Calcification.* The arteries were fixed in 10X neutral phosphate-buffered saline with 4% paraformaldehyde and embedded in paraffin blocks. Sections (4 μm) were stained for calcification using Von Kossa’s method as described previously^4^. Areas of calcification appeared as dark brown regions in the medial wall of the artery.

***Analysis***

Text data represented as mean±SD, unless otherwise indicated. The threshold for significance was a p-value <0.05. All statistical tests and graph generation were done on Graph Pad Prism (Version 8.4). Statistics performed are outlined in detail in figure captions and table footnotes.

**References for Complete Methods (all references also included in the full manuscript):**

1.	Shobeiri N, Pang J, Adams MA, Holden RM: Cardiovascular disease in an adenine-induced model of chronic kidney disease: the temporal link between vascular calcification and haemodynamic consequences. *J. Hypertens.* 31: 160–168, 2013

2. Kanagasabapathy DAS. WHO: Guidelines on Standard Operating Procedures for Clinical Chemistry. 2000:113.

3.	Heresztyn T, Nicholson BC: A colorimetric protein phosphatase inhibition assay for the determination of cyanobacterial peptide hepatotoxins based on the dephosphorylation of phosvitin by recombinant protein phosphatase 1. *Environ. Toxicol.* 16: 242–252, 2001

4. Proudfoot Diane, Skepper Jeremy N., Shanahan Catherine M. & Weissberg Peter L: Calcification of Human Vascular Cells In Vitro Is Correlated With High Levels of Matrix Gla Protein and Low Levels of Osteopontin Expression. *Arterioscler. Thromb. Vasc. Biol.* 18, 379–388, 1998.

**Appendix 1: Collected Tissues**

*Arteries:*

Aortic Arch

Proximal Thoracic Aorta

Distal Thoracic Aorta

Proximal Abdominal Aorta

Distal Abdominal Aorta

Carotid (L and R)

Celiac

Superior Mesenteric

Renal (L and R)

Iliac (L and R)

Pudendal (L and R)

*Veins:*

Vena Cava

Jugular (L and R)

*Bone:*

Lumbar Spine Vertebrae

Pelvic

Skull

Tibia

*Cardiac Muscle:*

Apex

Atria

Ventricle (L and R)

*Skeletal Muscle:*

Abdominal

Quadriceps

*Kidney:*

Medulla (L and R)

Cortex (L and R)

*Fat:*

Subcutaneous

Epididymal

*Miscellaneous:*

Skin

Testes

Bulbus

Lung (L and R)

Liver

Pancreas

Spleen

Mesentery
